# Theta-nested gamma oscillations in next generation neural mass models

**DOI:** 10.1101/2020.03.05.979021

**Authors:** Marco Segneri, Hongjie Bi, Simona Olmi, Alessandro Torcini

## Abstract

Theta-nested gamma oscillations have been reported in many areas of the brain and are believed to represent a fundamental mechanism to transfer information across spatial and temporal scales. In a series of recent experiments *in vitro* it has been possible to replicate with an optogenetic theta frequency stimulation several features of cross-frequency coupling (CFC) among theta and gamma rhythms observed in behaving animals. In order to reproduce the main findings of these experiments we have considered a new class of neural mass models able to reproduce exactly the macroscopic dynamics of spiking neural networks. In this framework, we have examined two set-ups able to support collective gamma oscillations: namely, the pyramidal interneuronal network gamma (PING) and the interneuronal network gamma (ING). In both set-ups we observe the emergence of theta-nested gamma oscillations by driving the system with a sinusoidal theta-forcing in proximity of a Hopf bifurcation. These mixed rhythms display always phase amplitude coupling. However two different types of nested oscillations can be identified: one characterized by a perfect phase locking between theta and gamma rhythms, corresponding to an overall periodic behaviour; another one where the locking is imperfect and the dynamics is quasi-periodic or even chaotic. From our analysis it emerges that the locked states are more frequent in the ING set-up. In agreement with the experiments, we find theta-nested gamma oscillations for forcing frequencies in the range [1:10] Hz, whose amplitudes grow proportionally to the forcing one and which are clearly modulated by the theta phase. Furthermore, analogously to the experiments, the gamma power and the frequency of the gamma-power peak increase with the forcing amplitude. At variance with experimental findings, the gamma-power peak does not shift to higher frequencies by increasing the theta frequency. This effect can be obtained, in or model, only by incrementing, at the same time, also the noise or the forcing amplitude. On the basis of our analysis both the PING and ING mechanisms give rise to theta-nested gamma oscillations with almost identical features.

## 1 INTRODUCTION

Oscillations in the brain, reflecting the underlying dynamics of neural populations, have been measured over a broad frequency range (Buzsaki, 2006). Particularly studied are *γ*-rhythms (30-120 Hz), due to their ubiquitous presence in many regions of the brain, irrespectively of the species (Buzsáki and Wang, 2012), and for their relevance for cognitive tasks (Fries et al., 2007) and neuronal diseases (Uhlhaas and Singer, 2006; Williams and Boksa, 2010).

Inhibitory networks have been shown to represent a fundamental ingredient for the emergence of *γ*-oscillations (Bartos et al., 2007; Buzsáki and Wang, 2012). Indeed, inhibition is at the basis of the two mainly reported mechanisms: pyramidal interneuronal network gamma (PING) and interneuronal network gamma (ING) (Tiesinga and Sejnowski, 2009). The ING mechanism is observable in purely inhibitory networks in presence of few ingredients: recurrent connections, a time scale associated to the synaptic *GABA_A_* receptors and an excitatory drive sufficiently strong to lead the neurons supra-threshold (Buzsáki and Wang, 2012). The collective oscillations (COs) emerge when a sufficient number of neurons begins to fire within a short time window and generate almost synchronous inhibitory post-synaptic potentials (IPSPs) in the post-synaptic interneurons. The inhibited neurons fire again when the IPSPs have sufficiently decayed and the cycle will repeat. Thus, the main ingredients dictating the frequency of the COs in the ING set-up are: the kinetics of the IPSPs and the excitatory drive (Whittington et al., 1995). On the othe hand the PING mechanism is related to the presence of an excitatory and an inhibitory population, then COs emerge whenever the drive on the excitatory neurons is sufficiently strong to induce an almost synchronous excitatory volley that in turn elicits an inhibitory one. The period of the COs is thus determined by the recovery time of the pyramidal neurons from the stimulus received from the inhibitory population (Wilson and Cowan, 1972). A peculiarity of this mechanism, observed both *in vivo* and *in vitro* experiments, is that there is a delay between the firing of the pyramidal cells and the interneuronal burst (Buzsáki and Wang, 2012).

Gamma oscillations are usually modulated by theta oscillations in several part of the brain, with theta frequencies corresponding to 4-12 Hz in rodents and to 1-4 Hz in humans. Specific examples have been reported for the hippocampus of rodents in behaving animals and during rapid eye movement (REM) sleep (Lisman, 2005; Colgin et al., 2009; Belluscio et al., 2012; Pernía-Andrade and Jonas, 2014; Colgin, 2015), for the visual cortex in alert monkeys (Whittingstall and Logothetis, 2009), for the neocortex in humans (Canolty et al., 2006) etc. This is an example of a more general mechanism of cross-frequency coupling (CFC) between a low and a high frequency rhythm which is believed to have a functional role in the brain (Canolty and Knight, 2010). In particular, low frequency rhythms (as the *θ* one) are usually involving broad brain regions and are entrained to external inputs and/or to cognitive events; while the high frequency activity (e.g. the *γ*-rhythm) reflects local computation activity. Thus CFC can represent an effective mechanism to transfer information across spatial and temporal scales (Canolty and Knight, 2010; Lisman and Jensen, 2013). Four different types of CFC of interest for electrophysiology has been listed in (Jensen and Colgin, 2007): phase-phase, phase-frequency, phase-amplitude and amplitude-amplitude couplings (PPC,PFC,PAC and AAC). Two more types of CFCs have been later added as emerging from the analysis of coupled nonlinear oscillators (Witte et al., 2008) and coupled neural mass models (Chehelcheraghi et al., 2017): frequency-frequency and amplitude-frequency coupling (FFC and AFC). In this paper we will consider *θ*-nested *γ* oscillations, and in this context we will analyze PPC, PFC and PAC between *θ* and *γ* rhythms. The most studied CFC mechanism is the PAC, which corresponds to the modification of the amplitude (or power) of *γ*-waves induced by the phase of the *θ*-oscillations. This mechanism has been reported in the primary visual cortex of anaesthetized macaques subject to naturalistic visual stimulation (Mazzoni et al., 2011), as well as during the formation of new episodic memories in the human hippocampus (Lega et al., 2016). As discussed in (Jensen and Colgin, 2007), *θ* phase can often modulate both amplitude (PAC) and frequency (PFC) of the *γ* oscillations, therefore these two mechanisms can occur at the same time. PPC, which refers to n:m phase locking between *γ* and *θ* phase oscillations (Tass et al., 1998), has been identified in the rodent hippocampus during maze exploration (Belluscio et al., 2012).

Out study is mostly motivated by recent optogenetic experiments revealing PAC in areas CA1 and CA3 of the hippocampus and in the medial enthorinal cortex (MEC) (Akam et al., 2012; Pastoll et al., 2013; Butler et al., 2016, 2018). These experiments have shown that a sinuoidal optogenetic stimulation at *θ*-frequency of the circuits *in vitro* is able to reproduce several features of *θ*-nested *γ* oscillations, usually observed in behaving rats (Bragin et al., 1995). All these experiments suggest that inhibition has a key role in generating this cross-frequency rhythms; however both ING (Pastoll et al., 2013) and PING (Butler et al., 2016, 2018) mechanism has been invoked to explain locally generated *γ* oscillations.

PING and ING oscillation mechanism have been qualitatively reproduced by employing heuristic neural mass models (Wilson and Cowan, 1972; Gerstner et al., 2014). However these standard firing rate models do not properly describe the synchronization and desynchronizaton phenomena occurring in neural populations (Devalle et al., 2017; Coombes and Byrne, 2019). Recently a new generation of neural mass models has been designed, which are able to exactly reproduce the network dynamics of spiking neurons of class I, for any degree of synchronization among the neurons (Montbrió et al., 2015). In particular, for purely inhibitory networks, these mean-field models have been able to reproduce the emergence of COs, observed in the corresponding networks, without the inclusion of an extra time delay (Devalle et al., 2017), as well as the phenomenon of event related synchronisation and desynchronisation (Coombes and Byrne, 2019).

Our main aim is to understand how *θ*-nested *γ* oscillations can emerge when a PING or ING mechanism is responsible for the fast oscillations and which differences can be expected in the population dynamics in the two cases. Therefore we will consider the new class of neural mass models introduced by Montbrió et al. (Montbrió et al., 2015) in two configurations: namely, a purely inhibitory population (ING set-up) and two coupled excitatory-inhibitory populations (PING set-up). In both configurations we will examine the system response to an external sinusoidal *θ*-drive.

In particular, Section II is devoted to the introduction of different spiking network configurations of Quadratic Integrate-and-Fire (QIF) neurons able to generate *γ* COs via PING and ING mechanisms and to the introduction of their corresponding exact neural mass formulations. A detailed bifurcation analysis of the neural mass models for the PING and ING set-ups, in absence of any external forcing, is reported in Section III. The PAC mechanism is analysed and discussed in Section IV. Firstly, by considering different types of PAC states: namely, phase locked or unlocked. Secondly, by comparing our numerical results for PAC dynamics with experimental findings, reported in (Butler et al., 2016) and (Butler et al., 2018), for the CA1 region of the hippocampus under sinusoidal optogenetic stimulations. Finally, a discussion of our results and of their implications, as well as of possible future developments, will be presented in Section V. The results reported in the paper are mostly devoted to super-critical Hopf bifurcations, however a specific example of sub-critical Hopf leading to COs is discussed for the PING set-up in Appendix A. Further network configurations ensuring the emergence of COs via PING mechanisms are presented in Appendix B.

## 2 MODELS AND BIFURCATION ANALYSIS

### 2.1 Network Models

In this paper we want to compare the two principal mechanism at the basis of the emergence of collective oscillatory dynamics in neural networks: namely, the PING and ING mechanisms. Therefore we will consider QIF neurons in the two following set-ups: namely, an excitatory and an inhibitory population coupled via instantaneous synapses (PING configuration) or a single inhibitory population interacting via post-synaptic potentials (PSPs) with exponential profile (ING configuration). The corresponding network configurations are shown in Fig. 1. Moreover the neurons are assumed to be fully coupled. As we will show in the following, both these two configurations support the emergence of COs.

**Figure 1.**
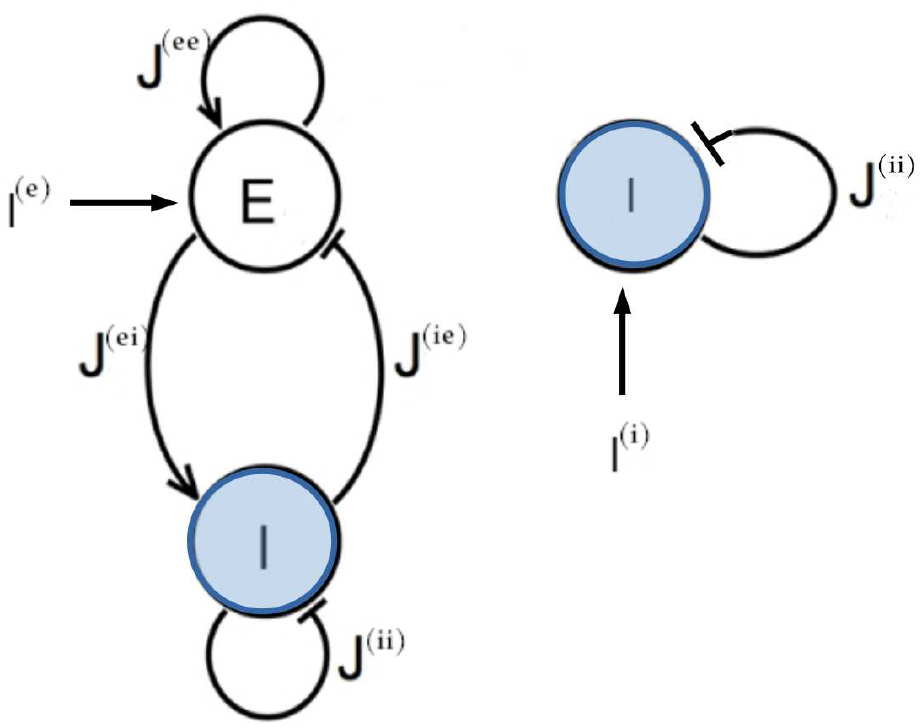
Network topologies. Two different network configurations have been investigated: on the left side, an excitatory population (E) and an inhibitory population (I) form a circuit that can generate oscillatory output (PING set-up); on the right side one inhibitory population (I) is coupled to itself with an inhibitory coupling (ING set-up). In both cases an external current *I*^(*l*)^ impinging one single population has been considered.

In particular, the dynamics of the membrane potentials of the QIF neurons in the PING configuration is given by

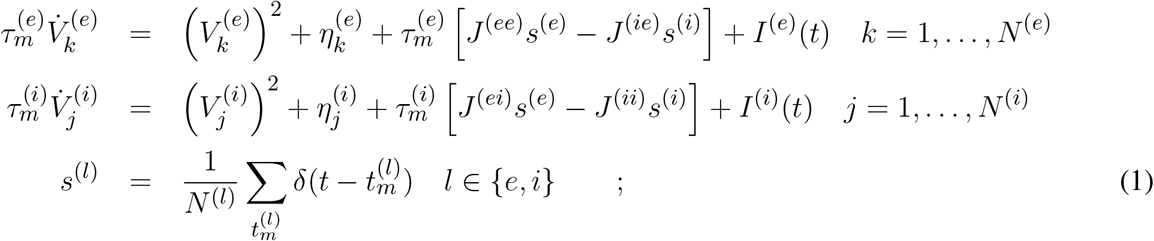

where the super-scripts *e* (*i*) denotes the excitatory (inhibitory) population, 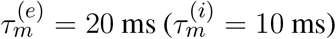 is the excitatory (inhibitory) membrane time constant, 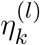 is the excitability of the *k*-th neuron of population *l*, *J*^(*ln*)^ is the strength of the synaptic coupling of population *l* acting on population *n*. The terms *I*^(*l*)^(*t*) represent a time dependent external current applied to the population *l*; usually we have considered the external drive to be applied to the excitatory population only, i.e. *I*^(*e*)^(*t*) ≠ 0 and *I*^(*i*)^(*t*) = 0. The synaptic field *s*^(*l*)^(*t*) is the linear super-position of all the pulses *p*(*t*) emitted in the past within the *l* population, being *p*(*t*) a *δ*-functions in the present case. Furthermore, since the neurons are fully coupled, each neuron will be subject to the same synaptic field (Olmi et al., 2010). The emission of the *m*-th spike in the network occurs at time 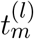 whenever the membrane potential of a generic neuron *j* reaches infinite, i.e. 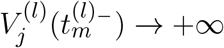, while the reset mechanism is modeled by setting 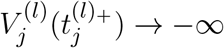, immediately after the spike emission.

The most part of our analysis of the PING set-up will be devoted to networks with self-activation only (i.e. where *J*^(*ii*)^ = 0), a configuration which is known to favour the emergence of collective oscillations (Wilson and Cowan, 1972; Kilpatrick, 2014; Onslow et al., 2014). However, as discussed in Appendix B, we have found that COs can arise in different PING set-ups: namely, in presence of self-inhibition only (i.e. with *J*^(*ii*)^ ≠ 0 and *J*^(*ee*)^ = 0) and in absence of both self-activation and inhibition (i.e. *J*^(*ee*)^ = *J*^(*ii*)^ = 0).

For what concerns the purely inhibitory network, the membrane potential dynamics of the *j*-th neuron is ruled by the following equations:

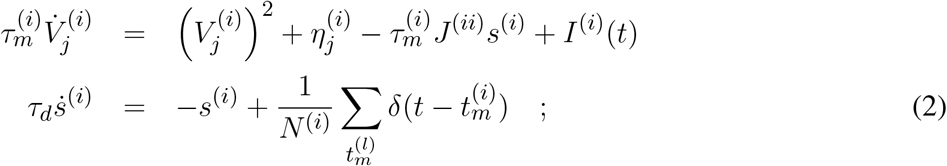

where 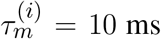. In this case the synaptic field *s*^(*i*)^(*t*) is the super-position of the exponential IPSPs 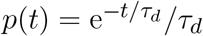 emitted in the past, where we set *τ_d_* = 10 ms.

For reasons that will be clarified in the next paragraph, we assume that the neuron excitabilities 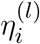 are randomly distributed according to a Lorentzian probability density function (PDF)

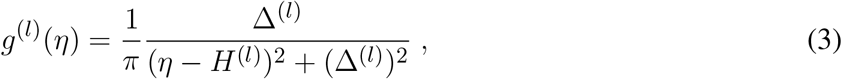

where *H*^(*l*)^ is the median and Δ^(*l*)^ is the half-width half-maximum (HWHM) of the PDF. Therefore each population will be composed by neurons supra- (with 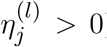) and sub-threshold (with 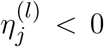), the percentage of one group with respect to the other being determined by the Lorentzian parameters. For the PING set-up we fix Δ^(*e*)^ = Δ^(*i*)^ = 1, while varying *H*^(*e*)^ and *H*^(*i*)^, while for the ING set-up we fix Δ^(*i*)^ = 0.3 and analyze the dynamics by varying *H*^(*i*)^.

The dynamical equations are integrated by employing a 4th order Runge-Kutta method in absence of noise with a time step *dt* = 0.002 ms (*dt* = 0.001 ms) for the PING (ING) set-up. Moreover, we define a threshold *V_p_* = 100 and a reset value *V_r_* = −100. Whenever the membrane potential *V_j_* of the *j*-th neuron overcomes *V_p_* at a time *t_p_*, it is reset to *V_r_* for a refractory period equal to 2/*V_j_*. At the same time the firing time is estimated as *t_p_* + 1/*V_j_*; for more details see (Montbrió et al., 2015). The membrane potentials are initialized from a random flat distribution defined over the range [−100 : 100], while the excitabilities are randomly chosen from the Lorentzian distribution (3).

In order to characterize the macroscopic dynamics we employ for instantaneous synapses the following indicators:

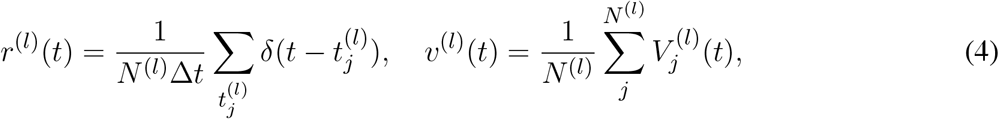

which represent the average population activity and the average membrane potential of a population *l*, respectively. In particular the average population activity of the *l*–network *r*^(*l*)^(*t*) is given by the number of spikes emitted in a time window Δ*t*, divided by the total number of neurons. For finite IPSPs we also consider the synaptic field *s*^(*l*)^(*t*). Furthermore, the emergence of COs in the dynamical evolution, corresponding to periodic motions of *r*^(*l*)^(*t*) and *υ*^(*l*)^(*t*), are characterized in terms of their frequencies *ν*(*l*).

We assume that the driving current, mimicking the *θ*-stimulation in the optogenetic experiments, is a purely sinusoidal excitatory current of the following form

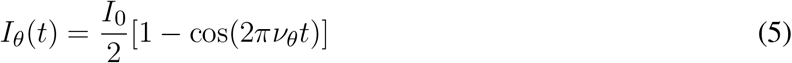

where *ν_θ_* is the forcing frequency, usually considered within the *θ*-range, i.e. *ν_θ_* ∈ [1 : 10] Hz. In this context a theta phase associated to the forcing field can be defined as *θ*(*t*) = *Mod*(2*πν*_*θ*_*t*, 2*π*). For the PING configuration we set *I*^(*e*)^(*t*) = *I_θ_*(*t*) and *I*^(*i*)^(*t*) ≡ 0 and for the ING set-up *I*^(*i*)^(*t*) = *I_θ_*(*t*).

### 2.2 Neural mass models

As already mentioned, an exact neural mass model has been derived by Montbrió et al. (Montbrió et al., 2015) for a fully coupled network of QIF neurons with instantaneous synapses and with Lorentzian distributed neuronal excitabilities. In this case the macroscopic neural dynamics of a population *l* is described by two collective variables: the mean field potential *υ*^(*l*)^(*t*) and the instantaneous firing rate *r*^(*l*)^(*t*). In this context, the neural mass model for two coupled *E* − *I* populations with instantaneous synapses, corresponding to the microscopic model reported in Eq. (1), can be written as

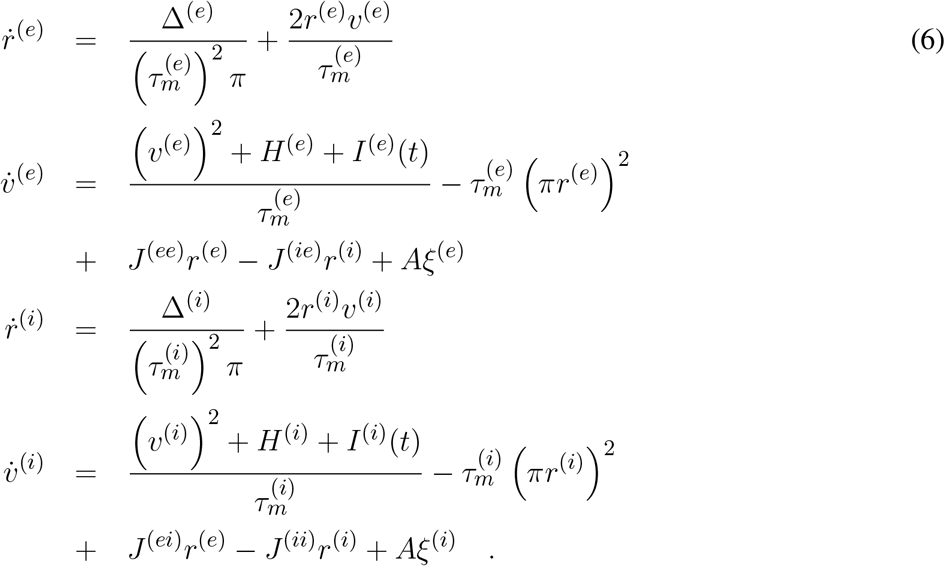

In the equations for the evolution of the average membrane potentials we have also inserted an additive noise term of amplitude *A*, employed in some of the analysis to mimic the many noise sources present in the brain dynamics. In particular the noise terms *ξ*^(*e*)^ and *ξ*^(*i*)^ are both *δ*-correlated and uniformly distributed in the interval [−1 : 1].

In case of finite synapses, the exact derivation of the corresponding neural mass model is still feasible for QIF neurons, but the macroscopic evolution now contains further equations describing the dynamics of the synaptic field characterizing the considered synapses (Devalle et al., 2017; Coombes and Byrne, 2019). In particular, for a single inhibitory population with exponential synapses, the corresponding neural mass model reads as:

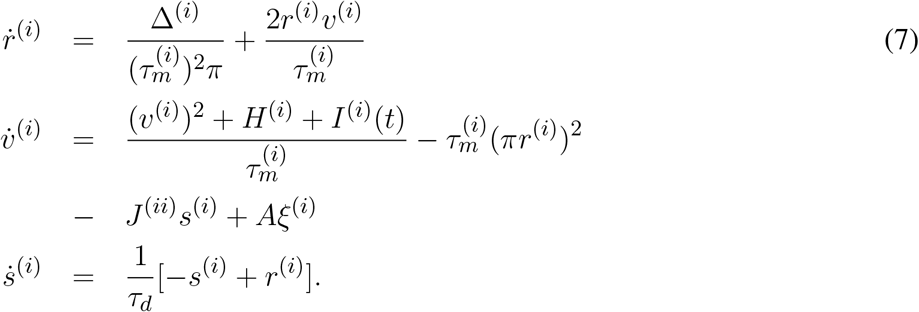

In the present case the equation for the average membrane potential contains, as already shown before in Eqs. (6), an additive noise term of amplitude *A*.

To analyse the stability of the macroscopic solutions of Eqs. (6) and (7), one should estimate the corresponding Lyapunov spectrum (Pikovsky and Politi, 2016). This can be done by considering the time evolution of the tangent vector, which for the PING set-up turns out to be four dimensional, i.e. *δ* = {*δr*^(*e*)^, *δυ*^(*e*)^, *δr*^(*i*)^*δυ*^(*i*)^}. The dynamics of the tangent vector is ruled by the linearization of the Eqs. (6), namely

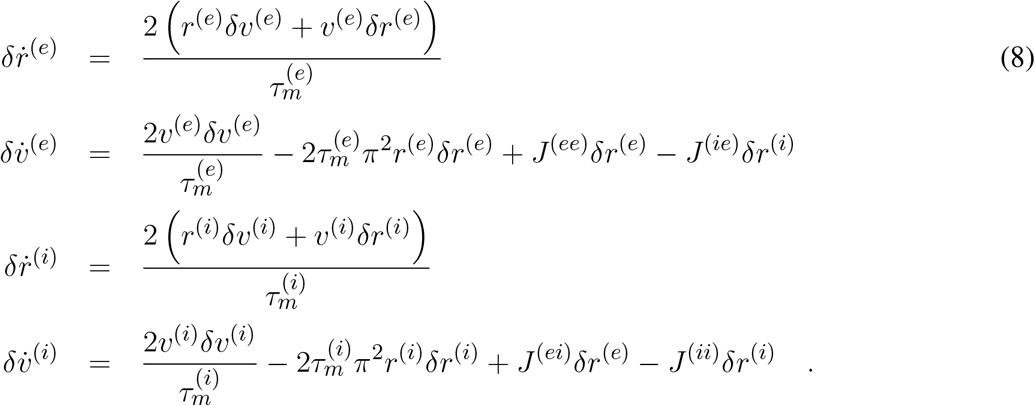

For the ING set-up the tangent vector is three dimensional, namely *δ* = {*δr*^(*i*)^, *δυ*^(*i*)^, *δs*^(*i*)^}, and its time evolution can be obtained by the linearization of the Eqs. (7), which reads as

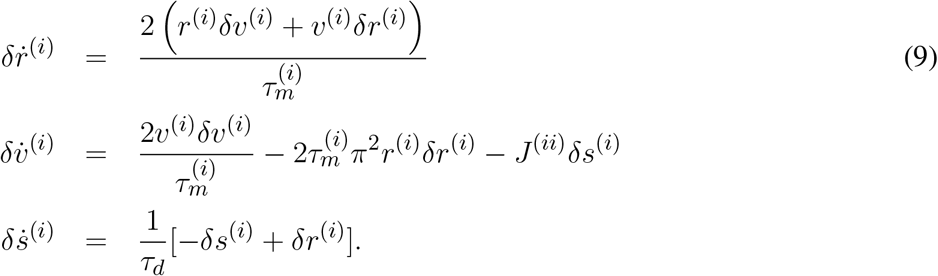

The LS is composed by 4 (3) Lyapunov exponents (LEs) {*λ_i_*} for the PING (ING) set-ups, which quantify the average growth rates of infinitesimal perturbations along the orthogonal manifolds. In details, LEs are estimated as follows

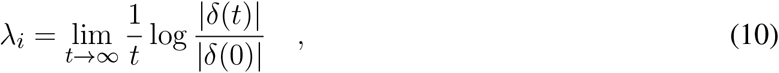

by employing the technique described in (Benettin et al., 1980) to maintain the tangent vectors orthonormal during the evolution. The autonomous system will be chaotic for *λ*_1_ > 0, while a periodic (quasi-perodic) dynamics will be characterized by *λ*_1_ = 0 (*λ*_1_ = *λ*_2_ = 0) and a fixed point by *λ*_1_ < 0. In a non-autonomous system in presence of an external forcing, one Lyapunov exponent will be necessarily zero, therefore a periodic behaviour corresponds to *λ*_1_ < 0 while a quasi-periodic one to *λ*_1_ = 0 (Pikovsky and Politi, 2016).

In absence of noise, neural mass models have been directly integrated by employing a Runge-Kutta 4th order integration scheme, while in presence of additive noise with a Heun scheme. In both cases the employed time step corresponds to *dt* = 0.01 ms. In order to estimate the Lyapunov spectra we have integrated the direct and tangent space evolution with a Runge-Kutta 4th order integration scheme with *dt* = 0.001 ms, for a duration of 200 s, after discarding a transient of 10 s.

Besides LEs, in order to characterize the macroscopic dynamics of the model, we have estimated the frequency power spectra 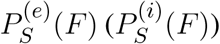 of the mean excitatory (inhibitory) membrane potential *υ*^(*e*)^(*t*) (*υ*^(*i*)^(*t*)) for the PING (ING) set-up. The power spectra have been obtained calculating the temporal Fourier transform of the mean membrane potentials sampled at time intervals of 2 ms. In the deterministic (noisy) case, time traces composed of 2048 (1024) consecutive intervals have been considered to estimate the spectra, which are obtained at a a frequency resolution of Δ*F* = 0.244 Hz (Δ*F* = 0.488 Hz). Finally, the power spectra have been averaged over 12 (488) independent realizations for the deterministic (noisy) dynamics. To compare our numerical results with the experimental ones reported in (Butler et al., 2016), as a measure of the power of the *γ* oscillations, we have estimated the area of the power spectrum *P_γ_* in an interval ±15 Hz around the main peak position *F_r_* of the corresponding power spectrum.

## 3 DYNAMICS IN ABSENCE OF FORCING

Due to the low dimensionality of the neural mass models we have been able to obtain the corresponding bifurcation diagrams by employing the software MATCONT developed for orbit continuation (Govaerts et al., 2006).

In particular, we have derived the bifurcation diagrams in absence of forcing (*I*^(*e*)^ = *I*^(*i*)^ ≡ 0) as a function of the median *H*^(*e*)^ and *H*^(*i*)^ (*H*^(*i*)^) of the excitability distributions for the PING (ING) configuration. In general, we observe either asynchronous dynamics, corresponding to a stable fixed point (a focus) of the neural mass equations, or COs, corresponding to stable limit cycles for the same set of equations.

### 3.1 PING set-up

For the excitatory-inhibitory set-up, as already mentioned, we usually fix *H*^(*i*)^ = −5 and we vary *H*^(*e*)^. In this case the inhibitory neurons are mostly below threshold (apart a 6-7 % of them) and they can be driven supra-threshold from the activity of the excitatory population for sufficiently large values of *H*^(*e*)^. COs emerge when a sufficient number of neurons is supra-threshold, i.e. when *H*^(*e*)^ becomes sufficiently positive. Indeed, as shown in Fig. 2 (a), at negative or low values of *H*^(*e*)^ one has asynchronous dynamics where the neurons fire indipendently an no collective behaviour is observable (as an example see Fig. 2 (c)). By increasing *H*^(*e*)^ a supercritical Hopf bifurcation occurs at 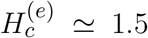 leading to the emergence of COs. The COs regime is characterized in the network by almost periodic population bursts, where the neurons in one population partially synchronize over a short time window of the order of few milliseconds. An example for *H*^(*e*)^ = 5 is shown in Fig. 2 (d), where one can observe two salient characteristics of the oscillatory dynamics. Firstly, the excitatory burst anticipates always the inhibitory one of a certain time interval *T_a_* (in this case *T_a_* ≃ 5 ms), as usually observed for the PING mechanism (Tiesinga and Sejnowski, 2009). Secondly, the bursts of the excitatory population have a temporal width (≃ 8 ms) which is two or three times larger than those of the inhibitory ones (≃ 2 − 3 ms). This is also due to the fact that a large part of the inhibitory neurons is sub-threshold, therefore most of them fire within a short time window, irrespectively of their excitabilities, due to the arrival of the synaptic stimulation from the excitatory population. Instead, the excitatory neurons, which are mostly supra-threshold, recover from silence, due to the inhibitory stimulation received during the inhibitory burst, over a wider time interval driven by their own excitabilities. It is evident that the COs frequency of the excitatory and inhibitory population coincide in this set-up.

**Figure 2.**
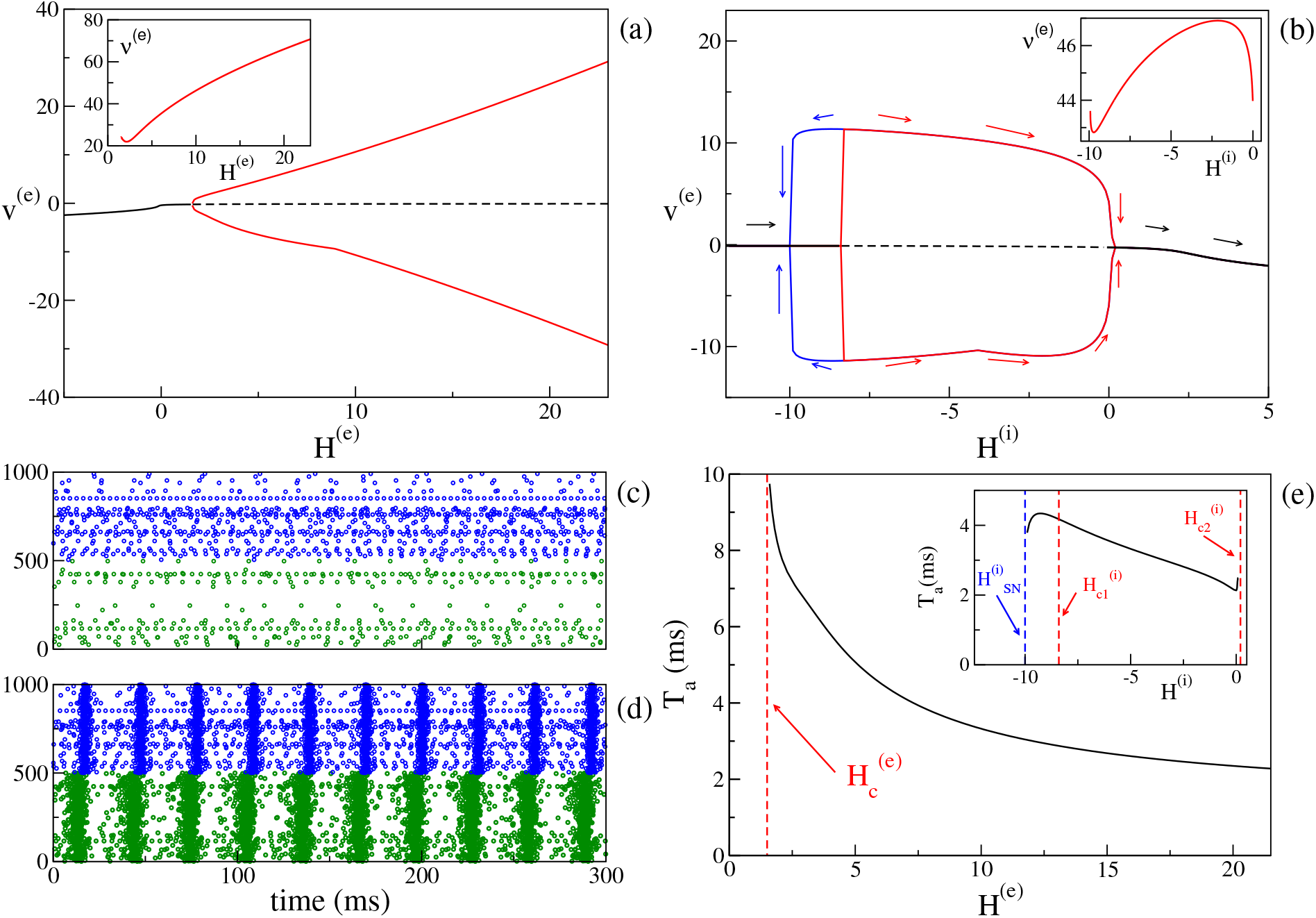
(PING set-up) (a) Bifurcation diagram of the average membrane potential *υ*^(*e*)^ as a function of *H*^(*e*)^ for *H*^(*i*)^ = −5.0. The black continuous (dashed) line identifies the stable (unstable) fixed point. The red lines denote the extrema of the limit cycles. The supercritical Hopf bifurcation occurs at 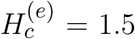. In the inset is reported the frequency *ν*^(*e*)^ of the COs versus *H*^(*e*)^. (b) Bifurcation diagram of the average membrane potential *υ*^(*e*)^ versus *H*^(*i*)^ for *H*^(*e*)^ = 10. The Hopf bifurcations are located at 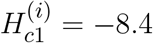 and 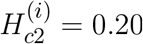, while the saddle-node of limit cycles at 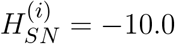. In the inset is reported the frequency *ν*^(*i*)^ ≡ *ν*^(*e*)^ of the COs versus *H*^(*i*)^. (c-d) Raster plots of the excitatory (green dots) and inhibitory (blue dots) networks are calculated in correspondence of the stable fixed point *H*^(*e*)^ = −5.0 (c) and of the limit cycle *H*^(*e*)^ = +5.0 (d) for the case analyzed in (a). For a better visualization, the activity of only 500 neurons of each population is shown. (e) Delay *T_a_* as a function of *H*^(*e*)^. The red dashed line denotes 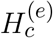. Here we have used the same parameters as in panel (a). In the inset is reported the dependence of *T*_*a*_ versus *H*^(*i*)^ for the parameters in panel (b). Other parameters of the system: *J*^(*ee*)^ = 8, *J*^(*ie*)^ = *J*^(*ei*)^ = 10, *J*^(*ii*)^ = 0 and sizes of the networks *N*^(*e*)^ = 5000, *N*^(*i*)^ = 5000.

Moreover it is important to investigate the bifurcation diagram of the system at fixed median excitatory drive by varying *H*^(*i*)^. The corresponding bifurcation diagram is reported in Fig. 2 (b) for *H*^(*e*)^ = 10. By increasing *H*^(*i*)^ COs emerge from the asynchronous state via a sub-critical Hopf bifurcation at 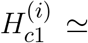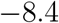 and they disappear via a super-critical Hopf at 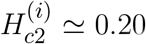. Since the first transition is hysteretical, COs disappear via a saddle-node of the limit cycles at a value 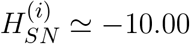 lower than 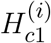. Indeed, in the interval 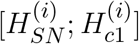 we have the coexistence of a stable focus with a stable limit cycle. In summary, COs are clearly observable as long as *H*^(*i*)^ is negative or sufficiently small. If the inhibitory neurons become mostly supra-threshold, this destroys the collective behaviour associated to the PING mechanism.

It is worth noticing that the frequencies of the COs are in the *γ*-range, namely *ν*^(*e*)^ ∈ [22 : 71] Hz (as shown in the inset of Fig. 2 (a)): in this set-up the maximal achievable frequency ≃ 100 Hz, since the decay time of inhibition is dictated by 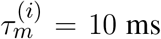 (Tiesinga and Sejnowski, 2009). On the other hand, the influence of *H*^(*i*)^ on the frequency of the COs is quite limited. As shown in the inset of Fig. 2 (b) for a specific case corresponding to *H*^(*e*)^ = 10.0, *ν*^(*i*)^ ≡ *ν*^(*e*)^ varies of few Hz (namely, from 42.8 to 46.9 Hz) by varying *H*^(*i*)^ of an order of magnitude.

For what concerns the delay *T_a_* between the excitatory and inhibitory bursts, we observe a decrease of *T_a_* with the increase of the excitatory drive *H*^(*e*)^, from *T_a_* ≃ 10 ms at the Hopf bifurcation, towards 2 ms for large *H*^(*e*)^ value, see Fig. 2 (e). The largest value of *T_a_* is of the order of 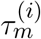. This can be explained by the fact that the excitatory stimulations should reach the inhibitory population within a time interval (at most) of 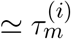 to be able to sum up in an effective manner and to ignite the inhibitory burst. As shown in the inset of Fig. 2 (e), the increase of *H*^(*i*)^ has in general the effect to reduce *T_a_*; this should be expected since for larger excitabilities (larger *H*^(*i*)^), the inhibitory neurons are faster in responding to the excitatory stimulations. However, this is not the case in proximity of the saddle-node bifurcation at 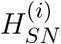 and for positive *H*^(*i*)^, where the effect is reversed and *T_a_* increases with *H*^(*i*)^.

For the PING set-up we can observe also sub-crtical Hopf bifurcations: a specific example is discussed in Appendix A in some details.

### 3.2 ING set-up

As shown in (Devalle et al., 2017), in order to observe COs in globally coupled inhibitory QIF networks and in the corresponding neural mass models, it is sufficient to include a finite synaptic time scale *τ_d_*. On the other hand, in sparse balanced QIF networks, COs are observable even for instantaneous synapses (di Volo and Torcini, 2018). Indeed, for the chosen set of parameters, by varying the median of the inhibitory excitabilities *H*^(*i*)^, we observe a super-critical Hopf bifurcation at 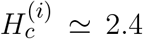, from an asynchronous state to a COs (see Fig. 3 (a)). Analogously to the PING set-up, the frequencies of the COs observable in the ING set-up are whithin the *γ*-range, namely *ν*^(*i*)^ ∈ [26 : 83] Hz. In particular, we observe an almost linear increases of *ν*^(*i*)^ with *H*^(*i*)^.

**Figure 3.**
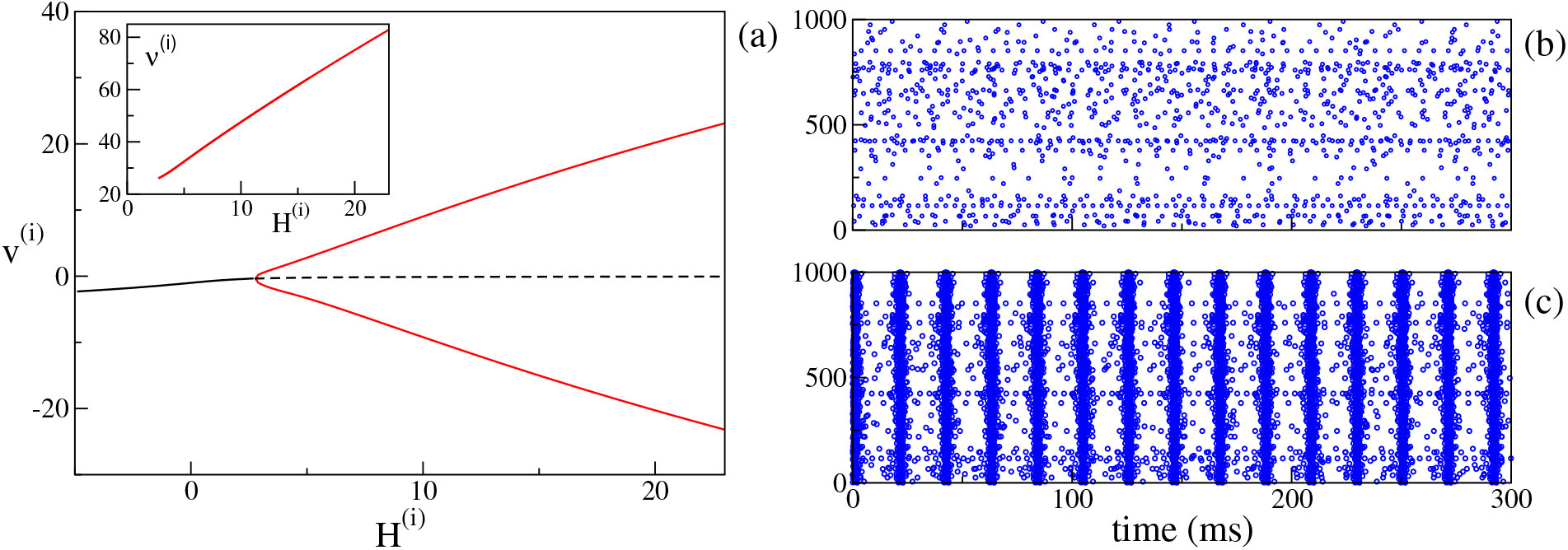
(ING set-up) (a) Bifurcation diagram of the average membrane potential *υ*^(*i*)^ as a function of *H*^(*i*)^. The black continuous (dashed) line identifies the stable (unstable) fixed point. The red lines denote the extrema of the limit cycles. The supercritical Hopf bifurcation occurs at 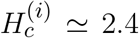. In the inset is reported the COs’ frequency *ν*^(*i*)^ of the inhibitory population as a function of *H*^(*i*)^. Right panels: raster plots of the inhibitory network (blue dots) are calculated in correspondence of the stable fixed point *H*^(*i*)^ = 0.0 (b) and of the limit cycle *H*^(*i*)^ = +10.0 (c). Only the firing activity of 1000 neurons is displayed. Parameters of the system: *J*^(*ii*)^ = 21.0, 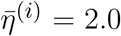, Δ^(*i*)^ = 0.3, 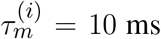, *τ*_*d*_ = 10.0 ms, *A* = 0. System size for the purely inhibitory network *N*^(*i*)^ = 10000.

Therefore, the PING and ING set-ups here considered are ideal candidates to analyse the influence of *θ*-forcing on *γ*-oscillatory populations, which represents the main focus of this paper. In particular, the response of the system to the excitatory *θ*-forcing current (5) can be interpreted in terms of the bifurcation diagrams for the model in absence of forcing shown, respectively, in Fig. 2 (a) for the PING set-up and in Fig. 3 (a) for the ING. The interpretation is possible due to the fact that the response of the system to the sinusoidal current (5) can be considered as almost-adiabatic, being the forcing frequencies *ν_θ_* ∈ [1 : 10] Hz definitely slower than those of the COs (*ν*^(*e*)^ and *ν*^(*i*)^), which lay in the *γ*-range.

## 4 DYNAMICS UNDER *θ*-FORCING

As a first step, we have verified that the reduced mean-field models are able to well reproduce the macroscopic evolution of the spiking network in both considered set-ups, under the external forcing (5).

In particular, we set the unforced systems in the asynchronous regime in proximity of a super-critical Hopf bifurcation, by choosing 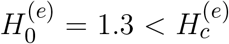 and 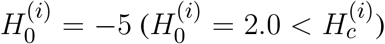 and considered a forcing term with frequency *ν_θ_* = 5 Hz and amplitude *I*_0_ = 10 (*I*_0_ = 9) for the PING (ING) set-up.

The comparisons, reported in Figs. 4 (a) and (c), reveal a very good agreement in both set-ups between the network and the neural mass simulations, for the mean membrane voltages and the instantaneous firing rates. Furthermore, in both cases, we clearly observe COs, whose amplitudes are modulated by the amplitude of *θ*-forcing term (5), suggesting that we are in presence of a Phase-Amplitude Coupling (PAC) mechanism (Hyafil et al., 2015). The corresponding spectrograms shown in Figs. 4 (b) and (d) reveal that the frequencies of the COs are in the *γ*-range with the maximum power localized around 50-60 Hz. Moreover the spectrograms indicate that the process is stationary and due to the external stimulation. The gamma oscillations repeat during each *θ*-cycles and they arrest when the external stimulation is stopped. The characteristics of these COs resemble *θ*-nested *γ* oscillations reported in many experiments for neural systems *in vitro* under optogenetic stimulation (Akam et al., 2012; Pastoll et al., 2013; Butler et al., 2016, 2018) as well as in behaving animals (Chrobak and Buzsáki, 1998).

**Figure 4.**
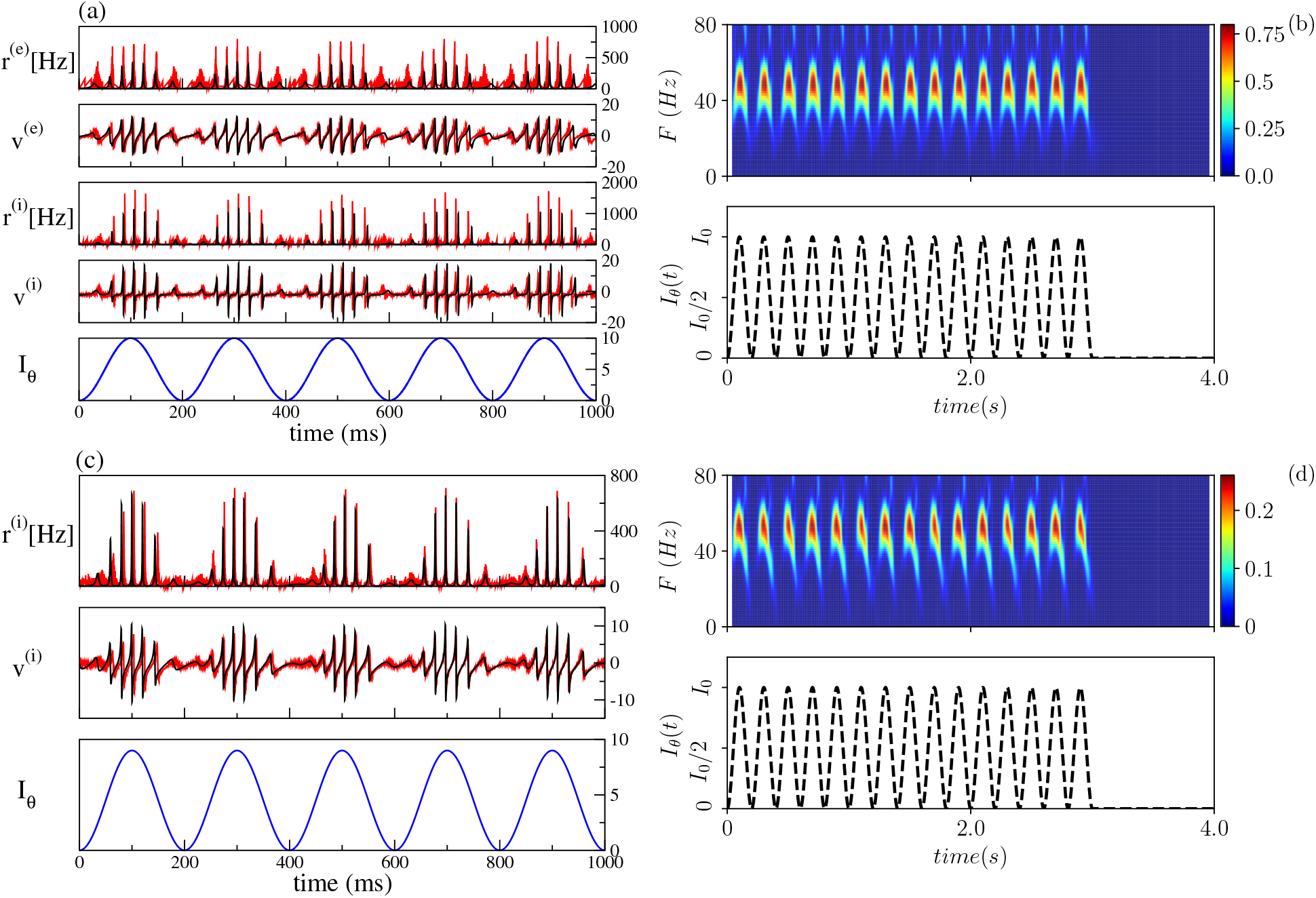
Theta-nested gamma oscillations (PING set-up) (a) From top to bottom: temporal traces of *r*^(*e*)^, *υ*^(*e*)^, *r*^(*i*)^, *υ*^(*i*)^, for the spiking network (red curves) and the neural mass model (black curves). *I*_*θ*_, reported in the bottom panel in blue, is the external current (5). For the neural mass model the average rates and membrane potentials are the solutions of Eqs. 6, while for the network they are calculated according to Eqs. 4. (b) Spectrogram of the mean membrane potential *υ*^(*e*)^ (top) as a function of the external forcing (bottom). The amplitude of the forcing is *I*_0_ = 10 and its frequency is *ν*_*θ*_ = 5 Hz. Parameters of the system: *J*^(*ee*)^ = 8, *J*^(*ie*)^ = *J*^(*ei*)^ = 10, *J*^(*ii*)^ = 0, 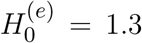, 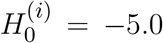, Δ^(*e*)^ = 1, 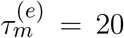, Δ^(*i*)^ = 1, 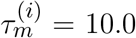, *A* = 0, network size *N*^(*e*)^ = *N*^(*i*)^ = 5000. The average firing rates are 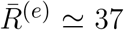 Hz, 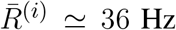. (**ING set-up**) (c) From top to bottom: temporal traces of *r*^(*i*)^, *υ*^(*i*)^ where the line colors have the same meaning as in panel (a). For the neural mass model, average rates and membrane potentials are solutions of Eqs. 7. (d) Spectrogram of the mean membrane potential *υ*^(*i*)^ (top) as a function of the external forcing (bottom). The amplitude of the forcing is *I*_0_ = 9 and its frequency is *ν*_*θ*_ = 5 Hz. Parameters of the system: *J*^(*ii*)^ = 21.0, 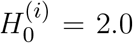, Δ^(*i*)^ = 0.3, 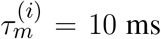, *τ*_*d*_ = 10.0 ms, *A* = 0, system size for the purely inhibitory network *N*^(*i*)^ = 10000. The corresponding average firing rate is 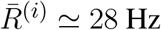.

### 4.1 Wavelet Analysis

To have a deeper insight on these dynamics we have estimated the continuous wavelet transform of the average membrane potential on each *θ*-cycle. As an example, we report in Fig. 5 the wavelet spectrogram of the mean potential within a single *θ*-cycle for the previously examined PING (panel (a)) and ING (panel (b)) set-ups. Indeed, from the comparison of panel (a) and (b) in Fig. 5, we pratically do not observe any difference: the system responds with COs in the range [40, 80] Hz and it exhibits alternating maxima and minima in the wavelet spectrogram as a function of the *θ*-phase. Similar results have been reported in (Butler et al., 2016) for the CA1 region of rat Hippocampus under optogenetic sinusoidal *θ* stimulation.

**Figure 5.**
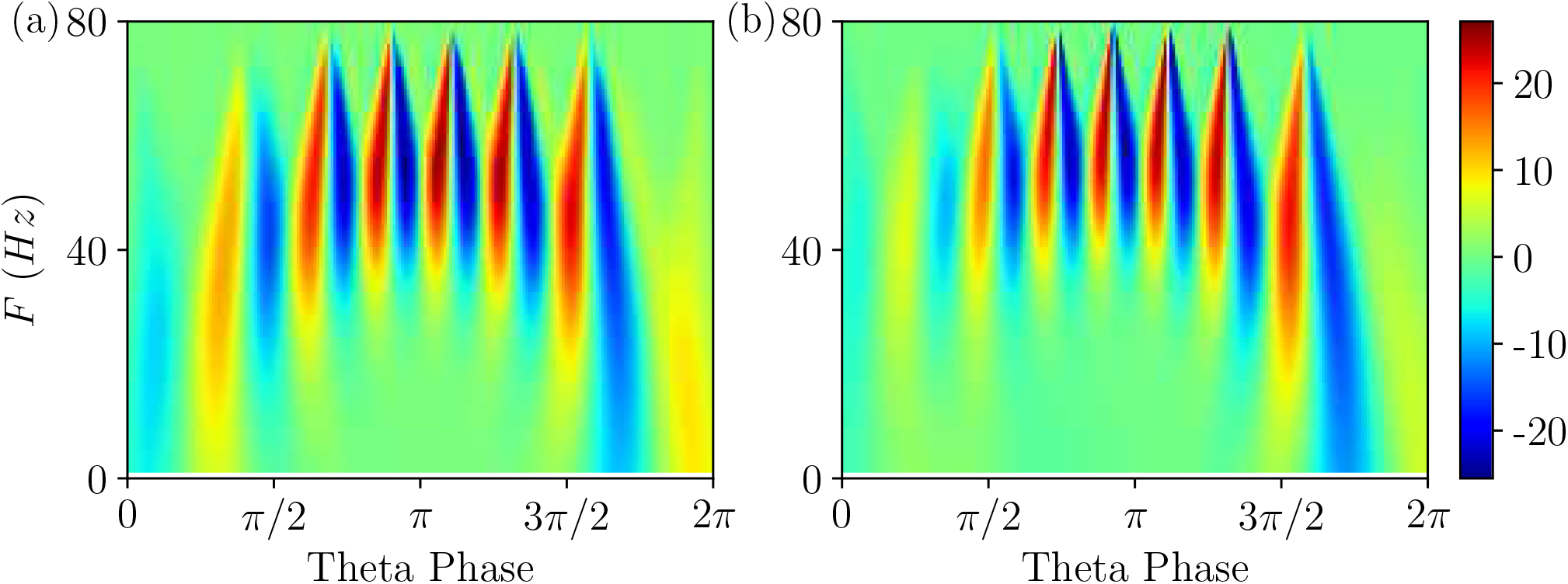
Wavelet Analysis. Continuous wavelet transform over a single *θ*-cycle of the mean membrane potentials *υ*^(*e*)^ and *υ*^(*i*)^ appearing in the neural mass models for PING (a) and ING (b) set-up, respectively. This analysis allows for accurate automated detection and extraction of gamma activity without the need for bandpass filtering. Parameters as in Fig. 4.

Differences among the two cases appear when one considers the wavelet spectrograms averaged over many *θ* periods: for the PING case the spectrogram remains unchanged, instead for the ING set-up the spectrogram smears out and it does not present anymore the clear oscillations reported in Fig. 5 (b). This difference indicates that in the PING case the observed pattern repeats exactly over each cycle: *γ* and *θ* oscillations are perfectly phase locked. This is not the case for the ING set-up: despite the PAC patterns appear quite similar in successive cycles, as shown in Fig. 4 (c), indeed they do not repeat exactly. From a point of view of nonlinear dynamics, the PING case would correspond to a perfectly periodic case, while the other case could be quasi-periodic or even chaotic. Therefore, we can observe PAC with associated phase locking or in absence of locking.

Furthermore, according to the data shown in Fig. 5, this can also represent an example of PFC, since COs with frequencies ≃ 40 Hz occur at small and large *θ*-phases, while in the middle range *π*/2 < *θ* < 3*π*/2 one observes similar oscillations with *F* ≃ 60 Hz.

### 4.2 Phase-Amplitude Locked and Unlocked States

To better examine the dynamical regimes emerging in our set-ups we have firstly estimated the maximal Lyapunov exponent *λ*_1_ associated to the neural mass models, for the same parameters considered in Fig. 4, over a wide range of forcing amplitudes, namely 0 ≤ *I*_0_ ≤ 20. From the results reported in Figs. 6 (a) and (b), it is clear that *λ*_1_ is always zero, apart from some limited intervals where it is negative. This means that the dynamics is usually quasi-periodic, apart from some Arnold tongues where there is perfect locking between the external forcing and the forced system.

**Figure 6.**
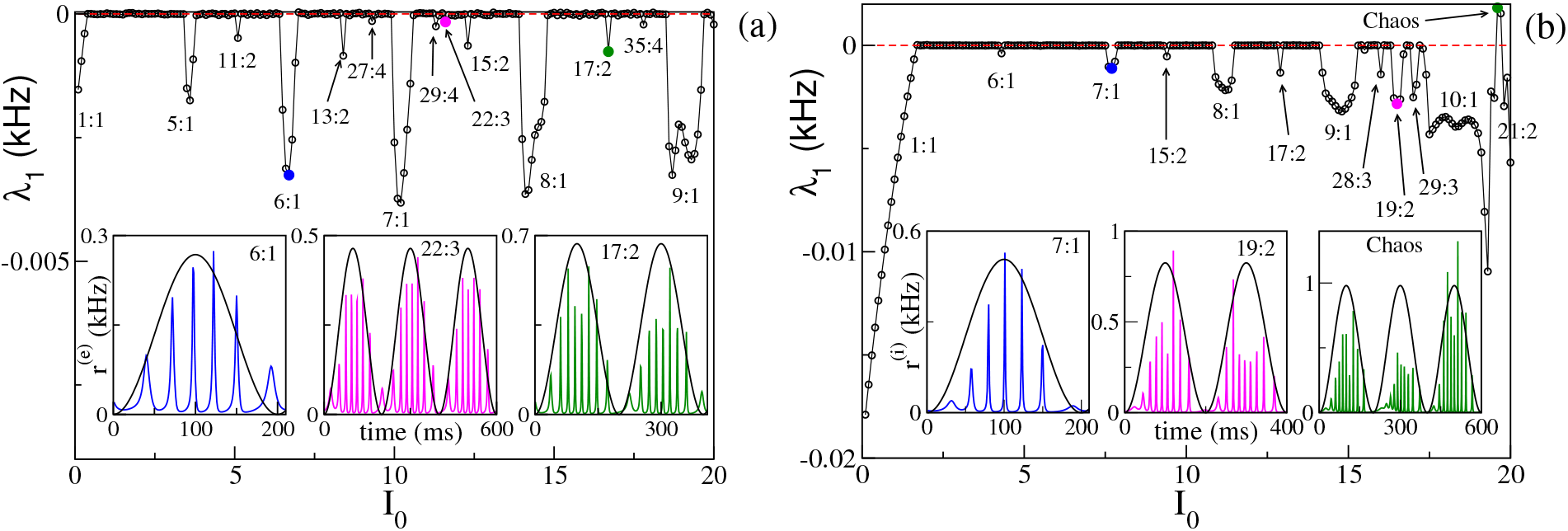
Maximal Lyapunov exponent. *λ*_1_ estimated for the neural mass model as a function of the forcing amplitude *I*_0_, for the PING (a) and ING (b) set-ups. In bothe cases the system is subject to a forcing frequency *ν_θ_* = 5 Hz. Insets in panel a (b) report the istantaneous firing rate *r*^(*e*)^(*t*) (*r*^(*i*)^(*t*)) versus time for the PING (ING) set-up respectively. The shown three cases are representative of the states identified by circles in the main panels. The color code is the same, i.e. the color used in the inset identifies the corresponding circle in the main panel. The black continuous lines in the inset corresponde to *I_θ_* in arbitrary units. Parameters as in Fig. 4.

We notice that for small amplitudes the forcing entrains the system in a 1 : 1 periodic locking, therefore the istantaneous firing rate displays one peak for each *θ*-period with the same frequency as the forcing *ν_θ_*. This locking is present in a wider region in the ING case (namely, *I*_0_ < 1.70) with respect to the PING set-up (namely, *I*_0_ < 0.40). More interesting locking regimes, where the forced populations oscillate in the *γ* range, emerge at larger *I*_0_. These locking regimes can be considered as *θ*-nested *γ* oscillations; mostly of them are of the type *m* : 1, with *m* ∈ [5 : 10], which means that, for each *θ*-period, the firing rate of the forced populations has *m* maxima (for specific examples see the insets of Fig. 6 (a) and (b)). In extremely narrow parameter intervals other, more complex, kinds of locking emerge of the type *m* : *n*, where exactly *m* maxima in the population activity appear for every *n θ*-oscillations. In the examined cases we have identified locked patterns with *n* up to four. Moreover, for the ING case, we have observed even a chaotic region (see Fig. 6 (b)), which emerges at quite large forcing amplitude *I*_0_ ≃ 19. On the basis of our analysis we cannot exclude that chaos could emerge also in the PING set-up, for sufficiently strong forcing.

Let us now focus on the *m* : 1 perfectly locked states with *m* > 1, which are worth investigating due to their relevance for *θ*-*γ* mixed oscillations as well as to their relative large frequency of occurence with respect to more complex *m* : *n* locked states. In particular, we have examined the response of the system to different forcing amplitudes *I*_0_ ∈ [0 : 20] and frequencies *ν_θ_* ∈ [1 : 10] Hz. The *m* : 1 locked oscillations are reported in Figs. 7 (a) and (b) and characterized by the number *m* of oscillations displayed within a single *θ*-cycle.

**Figure 7.**
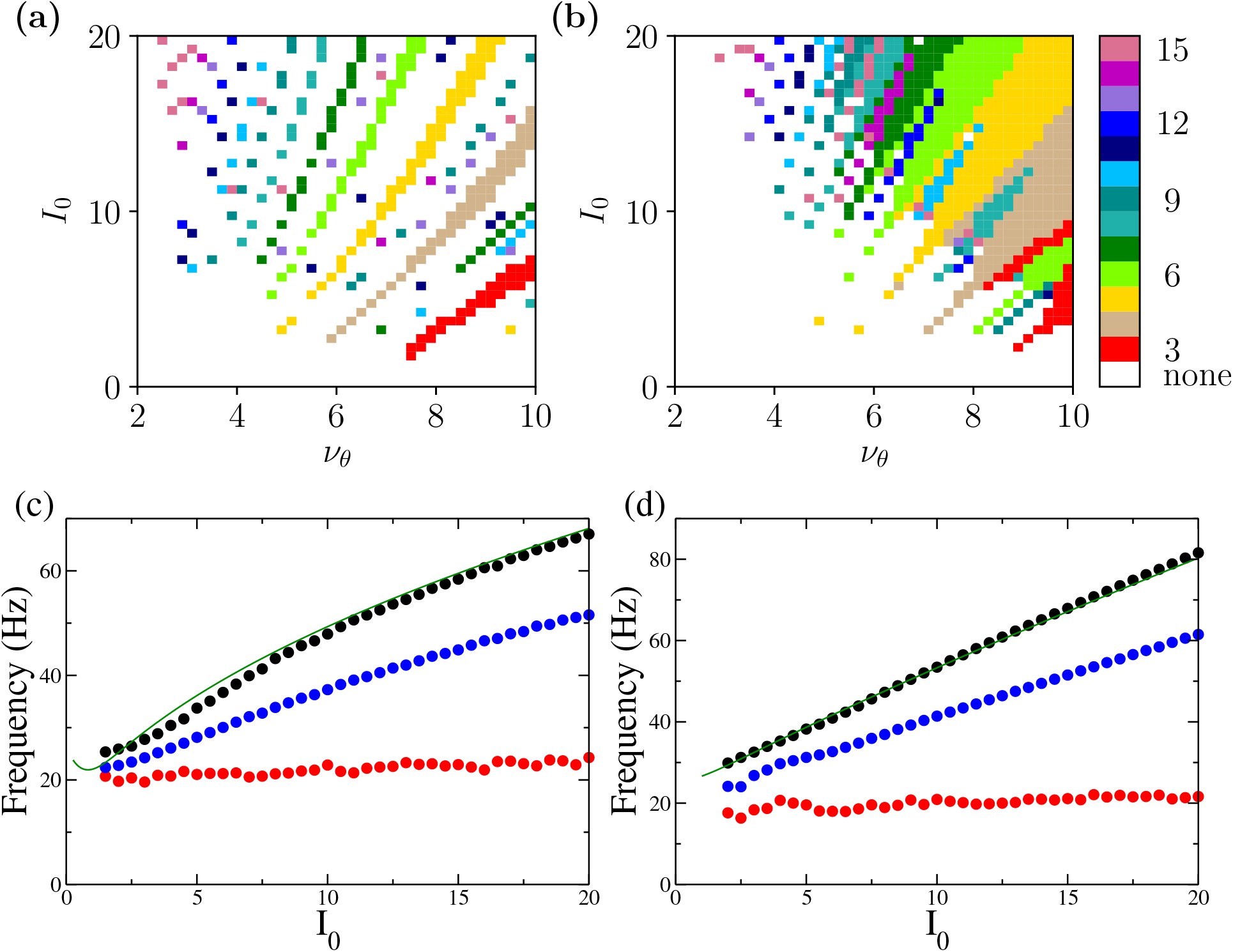
Phase locked *m* : 1 states. Locked states are displayed in panels (a) and (b) for the PING and ING set-ups, respectively. The color code identifies the locked states accordingly to *m*, from 3 to 15. (c,d) Minimal (red circles), average (blue circles) and maximal (black circles) frequencies of the COs as a function of the forcing amplitude *I*_0_ for PING (c) and ING (d) set-ups. These values are obtained by considering all possible *m* : 1 locked states corresponding to the examined *I*_0_. The frequencies *ν*^(*e*)^ (*ν*^(*i*)^) (green solid lines) of the COs obtained from the bifurcation analysis in the adiabatic set-up are reported as a function of 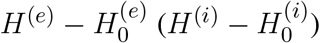 for the PING (ING). Parameters as in Fig. 4.

These locked states appear only for *ν_θ_* > 2 − 3 Hz. Moreover the states with equal *m* are arranged in stripes in the (*ν_θ_*, *I*_0_)-plane. Locked states in the PING configuration occur in separated stripes whose order *m* increases for increasing *I*_0_; in particular states with 3 ≤ *m* ≤ 10 are clearly identifiable. In the ING set-up, for sufficiently large *ν_θ_* and *I*_0_, we have a continuum of locked states, thus indicating that, for the ING set-up, phase locking to the forcing frequency is easier to achieve. In this case the order of occurrence of *m*-order states is not clearly related to the forcing amplitude; however locked states with order *m* and 2*m* are often nested within each other as shown in Fig. 7 (b).

To examine which frequencies are excited in these states we have measured, for each amplitude *I*_0_, the minimal, the maximal and the average frequency of the COs associated to *m* : 1 locked states over the whole range of examined forcing frequencies *ν_θ_*. These frequencies are reported in Figs. 7 (c) and (d). The analysis clearly reveals that the minimal CO frequency is essentially independent from *I*_0_ and its value is around 20 Hz, while the maximal and the average ones grow with *I*_0_. However all the frequencies stay within the *γ*-range for the examined forcing amplitudes.

To better understand the mechanism underlying the emergence of *θ*-nested *γ*-oscillations, we have reported in Figs. 7 (c) and (d) the COs’ frequencies *ν*^(*e*)^ (*ν*^(*i*)^) (green solid lines) obtained from the adiabatic bifurcation analysis of the neural mass models (these frequencies are also shown in the insets of Figs. 2 (a) and 3 (a)). The very good agreement between *ν*^(*e*)^ and *ν*^(*i*)^ and the maximal frequency measured for the locked states suggests that the nested COs are due to the crossing of the super-critical Hopf bifurcation during the periodic stimulation. In particular, during forcing, the maximal achievable *γ*-frequency is the one corresponding to the maximal stimulation current 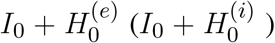 for the unforced PING (ING) set-ups. Furthermore, under sinusoidal forcing, the system spends a longer time in proximity of the maximal stimulation value, since it is a turning point. This explains why this frequency is always present in the response of the driven system for the considered locked states.

### 4.3 Comparison with Experimental Findings

In a series of recent optogenetic experiments on the mouse enthorinal-hippocampal system, have been reported clear evidences that phase-amplitude coupled *γ* rhythms can be generated locally in brain slices *ex vivo* in the CA1 region, as well as in the CA3 and MEC under sinusoidal *θ* stimulations (Akam et al., 2012; Pastoll et al., 2013; Butler et al., 2016, 2018). In particular, in (Butler et al., 2018) the authors reported evidences that, under theta-rhythmic activation of pyramidal neurons, the generation of the *γ* rhythms is due to a PING mechanism in all the three mentioned regions. However, due to the fact that pyramidal neurons are directly activated during experiments, their result cannot exclude that tonic activation of interneurons contributes to *θ*–*γ* oscillations *in vivo*. Furthermore, in (Pastoll et al., 2013) the authors affirm that in MEC *θ*-nested *γ* oscillations due to the optogenetic *θ* frequency drive, are generated by local feedback inhibition without recurrent excitation, therefore by a ING mechanism. In this Section we would try to reproduce some of the analyses reported in these experimental studies by employing both the PING and ING set-ups, in order to understand if these two set-ups give rise to different dynamical behaviours.

By following the analysis performed in (Butler et al., 2016, 2018), we considered the response of the two set-ups to forcing of different frequencies *ν_θ_* and amplitudes *I*_0_. The results reported in Fig. 8 reveal that the phenomenon of PAC is present for all the considered frequencies *ν_θ_* ∈ [1, 10] Hz and amplitudes *I*_0_ ∈ [1, 20] in both set-ups. Moreover, analogously to what reported in (Butler et al., 2016, 2018), the amplitude of the *γ* oscillations increases proportionally to *I*_0_ while the number of nested oscillations in each cycle increases for decreasing *ν_θ_*. On the basis of this comparison, the forced PING and ING set-ups display essentially the same dynamics.

**Figure 8.**
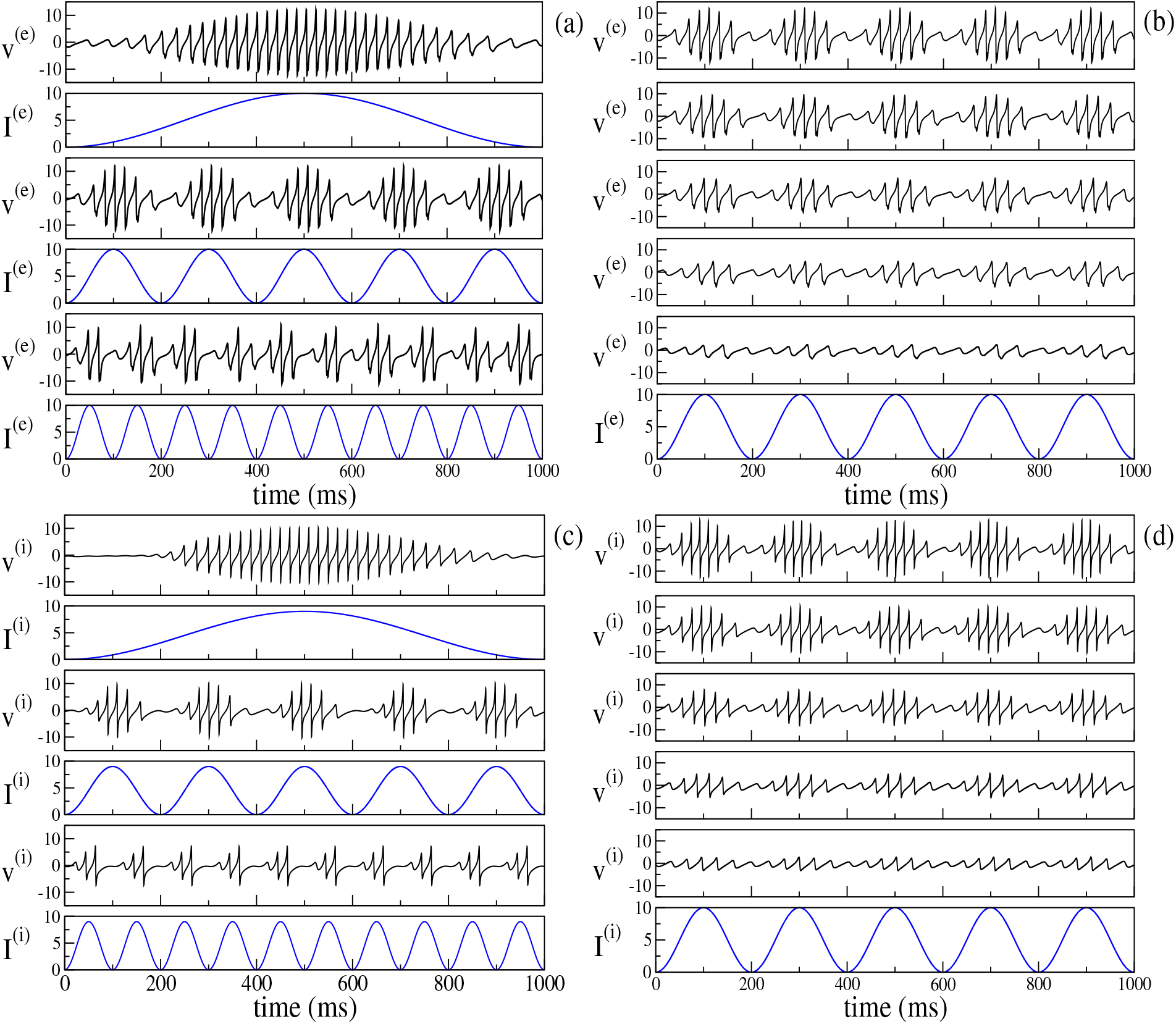
Theta-Nested Gamma COs for PING (a-b) and ING set-up (c-d) Left column: dependence of the mean membrane potential of the excitatory (inhibitory) population *υ*^(*e*)^ (*υ*^(*i*)^) on the frequency *ν_θ_* of the external forcing *I*^(*e*)^ = *I_θ_* (*I*^(*i*)^ = *I_θ_*) with *I*_0_ = 10 (*I*_0_ = 9) for the PING (ING) set-up. The current profiles (blue lines) are displayed immediately below the corresponding membrane potential evolution. From top to bottom, the frequency *ν_θ_* is 1 Hz, 5 Hz and 10 Hz. Right column: dependence of the mean membrane potential *υ*^(*e*)^ (*υ*^(*i*)^) on the amplitude *I*_0_ of the external current. Here the forcing frequency is kept constant at the value *ν_θ_* = 5*Hz*. The amplitude is changed from 20% of maximum (bottom) to 100% of maximum (top) in 20% increments, the maximum being given by *I*_0_ = 10. Other parameters as in Fig. 4.

To get a more detailed information about the dynamics in the two set-ups, we will now consider the features of the power spectra 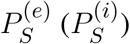 of the mean excitatory (inhibitory) potential for the PING (ING) set-up. These features are obtained for different forcing amplitudes and frequencies, somehow similarly to the analysis performed for the power spectra of the Local Field Potential (LFP) in (Butler et al., 2016, 2018).

Let us first consider, as an example of the obtained power spectra, the case corresponding to the PING set-up with a forcing characterized by *ν_θ_* = 5 Hz and amplitude *I*_0_ = 10, shown in Fig. 9 (a). In the spectrum we observe very well defined spectral lines located at frequencies which can be obtained as a linear combination of the forcing frequency *ν_θ_* = 5 Hz and of the frequency *F_r_* = 45 Hz. In particular *F_r_* is associated to the main peak and should correspond to the intrinsic frequency of the forced system. In the present case, the adiabatic bifurcation diagram reported in Fig. 2 (a) tells us that the maximal achievable COs’ frequency is 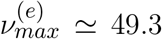, corresponding to 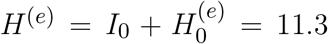. Indeed 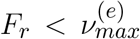 and this is due to the fact that the interaction with the forcing system can induce a locking phenomenon at a frequency that is exactly a multiple of *ν_θ_*, as it happens in the present case. However, in general, a spectrum as the one shown in Fig. 9 (a) is the emblem of a quasi-periodic motion characterized by two uncommensurate frequencies. This can be easily observable in most of the cases in our system, where *ν_θ_* and *F_r_* are usually uncommensurate.

**Figure 9.**
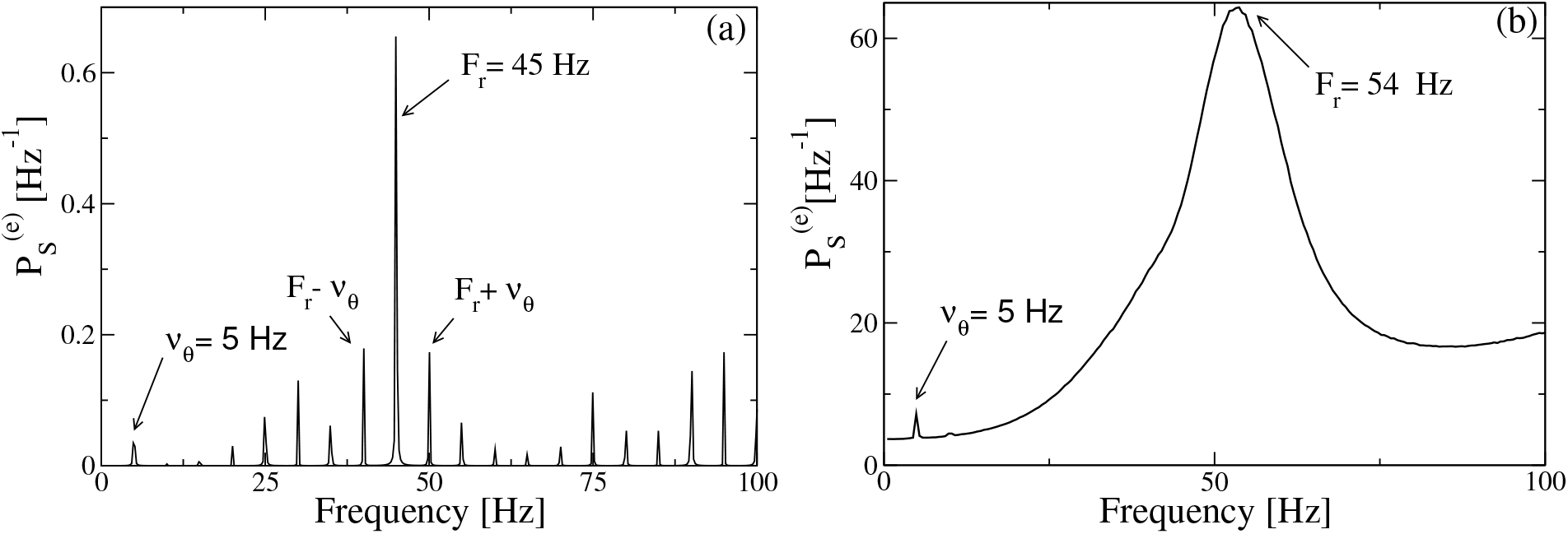
Power spectra for the PING set-up. Spectra 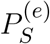 of the mean membrane potential *υ*^(*e*)^ estimated when the excitatory population is subject to an external drive with frequency *ν_θ_* = 5 Hz and amplitude *I*_0_ = 10, in absence of noise (a) and for additive noise with amplitude *A* = 1.4 (b). Other parameters as in Fig. 4.

The spectra obtained from optogenetic stimulation, reported in (Butler et al., 2016, 2018), do not resemble the one shown in Fig. 9 (a); indeed they present only two peaks: one corresponding to the stimulation frequency and one, quite broad, associated to the *γ* oscillations. We can expect that the difference is due to the multiple noise sources that are always present in an experimental analyis (and in particular for neurophysiological data), but that are absent in our model. Indeed, by considering the neural mass model for the PING set-up with additive noise on the membrane potentials of suitable amplitude, namely *A* = 1.4, we get a power spectrum resembling the experimental ones, as shown in Fig. 9 (b). The presence of noise induces the merging of the principal peaks in an unique broad one and the shift of the position of the main peak towards some larger values (namely, *F_r_* = 54 Hz in the present case) with respect to the fully deterministic case.

Let us now consider the power spectra obtained for different forcing frequencies *ν_θ_* ∈ [1 : 10] Hz in the *θ* range, in case of fixed forcing amplitude and in absence of noise. The position of the main and auxiliary peaks are shown in Fig. 10 (a) (Fig. 10 (c)) for the PING (ING) set-up and compared with the experimental results (red circles) obtained for the CA1 region of the hippocampus in (Butler et al., 2016). It is clear that, for both set-ups, the position of the main peak *F_r_* (green squares) has a value ≃ 50 Hz and it does not show any clear dependence on *ν_θ_*. This is in contrast with the experimental data, which reveals an increase proportional to *ν_θ_* from 49 Hz to 60 Hz. The same trend is displayed in our simulation from the subsidiary peak located at *F_r_* + *ν_θ_* (black stars), that somehow obviously increases with *ν_θ_*.

**Figure 10.**
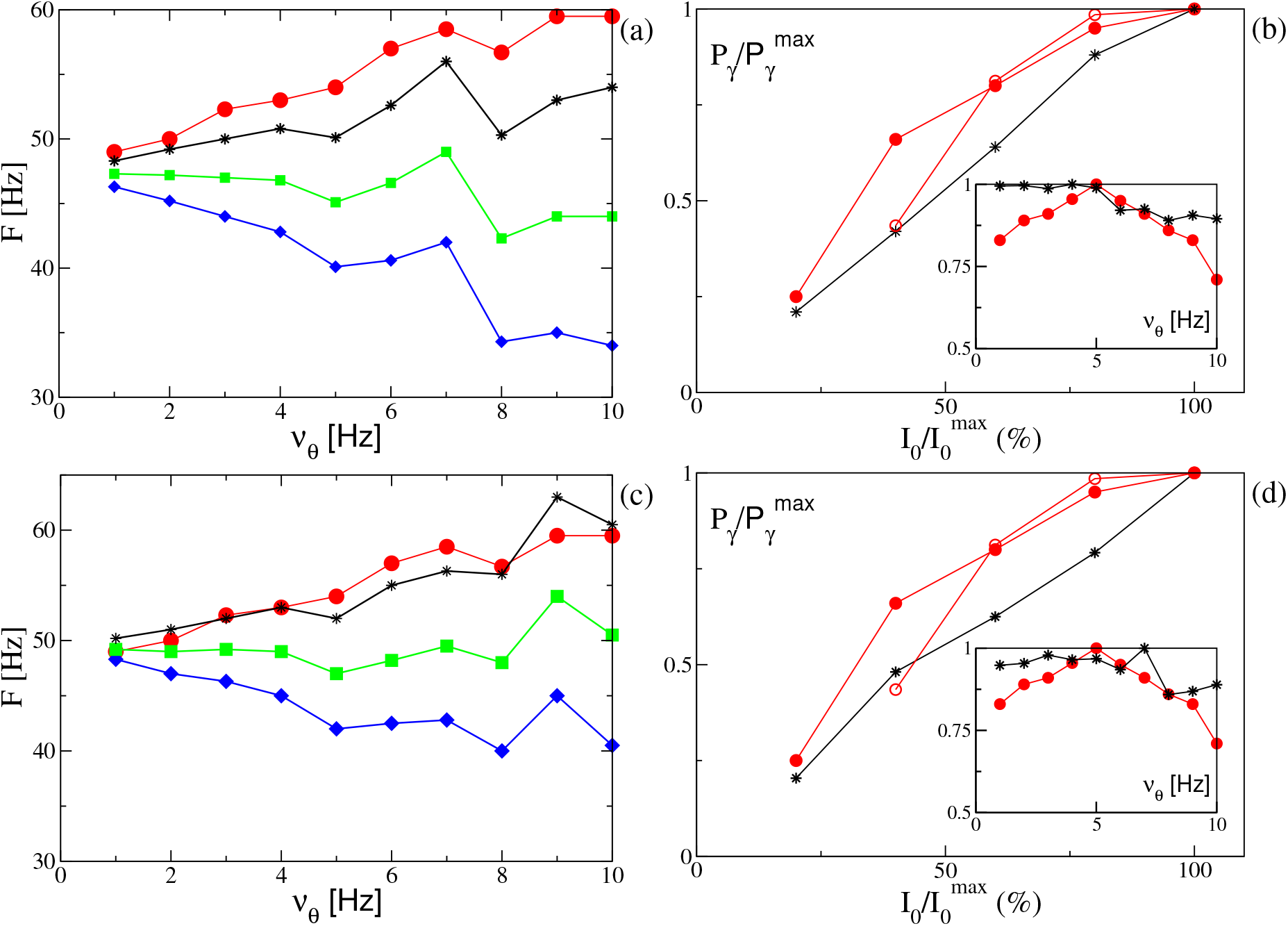
Power spectra features (PING set-up) (a) Frequencies of the peaks of the power spectrum 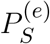 as a function of the stimulation frequency *ν_θ_*. Green squares correspond to the main peak frequency *F_r_*, while the black stars to *F_r_* + *ν_θ_* and the blue diamonds to *F_r_* − *ν_θ_*. The red circles are the experimental data extrapolated from Fig. 4C of (Butler et al., 2016). The amplitude of the forcing is *I*_0_ = 10. (b) Normalized power of the *γ* oscillations 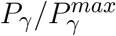 associated to the signal *υ*^(*e*)^ as a function of the amplitude stimulation, where we set 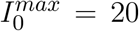 and the frequency of stimulation at *ν_θ_* = 5 Hz. In the inset we report the same quantity as a function of the frequency stimulation *ν_θ_* for *I*_0_ = 10. The black stars correspond to our simulations, while the red circles to experimental data extrapolated from Fig. 4E (Fig. 4B for the inset) of Ref. (Butler et al., 2016) (filled circles) and from Fig. 4C of Ref. (Butler et al., 2018) (empty circles). The other parameters are as in Fig. 4. (**ING set-up**) (c) Same as in panel (a) for the power spectrum 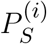 with *I*_0_ = 9. (d) Same as panel (b) for the signal *υ*^(*i*)^ with 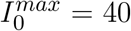. For the inset we set *I*_0_ = 9, other parameters as in Fig. 4.

Let us now consider the power of the *γ* oscillations *P_γ_* as defined in sub-section II B. As shown in the insets of Fig. 10 (b) and (d), this quantity remains essentially constant for low frequencies (namely, for *ν_θ_* ≤ 5 Hz in the PING and for *ν_θ_* ≤ 7 Hz in the ING), while it drops to smaller values at larger frequencies. On the other hand, the experimental results (red circles) reveal a similar decrease at frequencies *ν*_θ_ > 5 Hz, but they also reveal an increase at low frequencies, not present in our numerical data, thus suggesting a sort of resonance at 5 Hz. For what concerns the dependence of *P_γ_* on the forcing amplitude, we have fixed *ν_θ_* = 5 Hz and varied *I*_0_ in the range [4 : 10] ([8 : 20]) for the PING (ING) set-up. In both cases and analogously to experimental data, *P_γ_* increases proportionaly to *I*_0_, see Fig. 10 (b) and (d).

Our model is unable to reproduce, in both set-ups, in absence of noise and for fixed forcing amplitude *I*_0_, the steady increase of *F_r_* with *ν_θ_* reported in the experiments for the CA1 of mice in (Butler et al., 2016). Therefore, in order to cope with this problem, we will now investigate how a similar trend can emerge in our data. In particular, in the remaining part of the paper we consider noisy dynamics, to have a better match with experiments where it is unavoidable. In Fig. 11 (a) we report, for the PING set-up, the estimated power spectra for different noise levels, under constant external sinusoidal forcing. The effect of noise is to render the spectrum more flat and to shift the position of the peak in the *γ* range towards higher frequencies. As shown in the inset of Fig. 11 (a), the frequency *F_r_* is almost insensitive to the noise up to amplitudes *A* ≃ 1.0, then it increases steadily with *A* from ≃ 45 hz to ≃ 62 Hz. The effect of varying the forcing amplitude *I*_0_, for constant forcing frequency *ν_θ_* = 5 Hz and noise amplitude *A* = 1.4, is shown in Fig. 11 (b). In this case the amplitude increase of the forcing leads to more defined peaks in the *γ* range and to an almost linear increase with *I*_0_ of *F_r_*, as reported in the inset. In the same inset we have reported also the results related to two optogenetic experiments for the CA1 region of the mice hippocampus. In particular the data sets refer to two successive experiments performed by the same group and reported in (Butler et al., 2016, 2018). While in one experiment (red open circles) a clear increase of *F_r_* with the forcing amplitude is observable from 60 to 70 Hz, in the another one (red filled circles) the frequency remains almost constant around 45 − 50 Hz for a variation of *I*_0_ from 40 to 100 % of the maximal amplitude (Butler et al., 2018). From this comparison, we can affirm that our data catch the correct range of frequencies in both experiments and the dependence on the forcing amplitude reported in (Butler et al., 2018).

**Figure 11.**
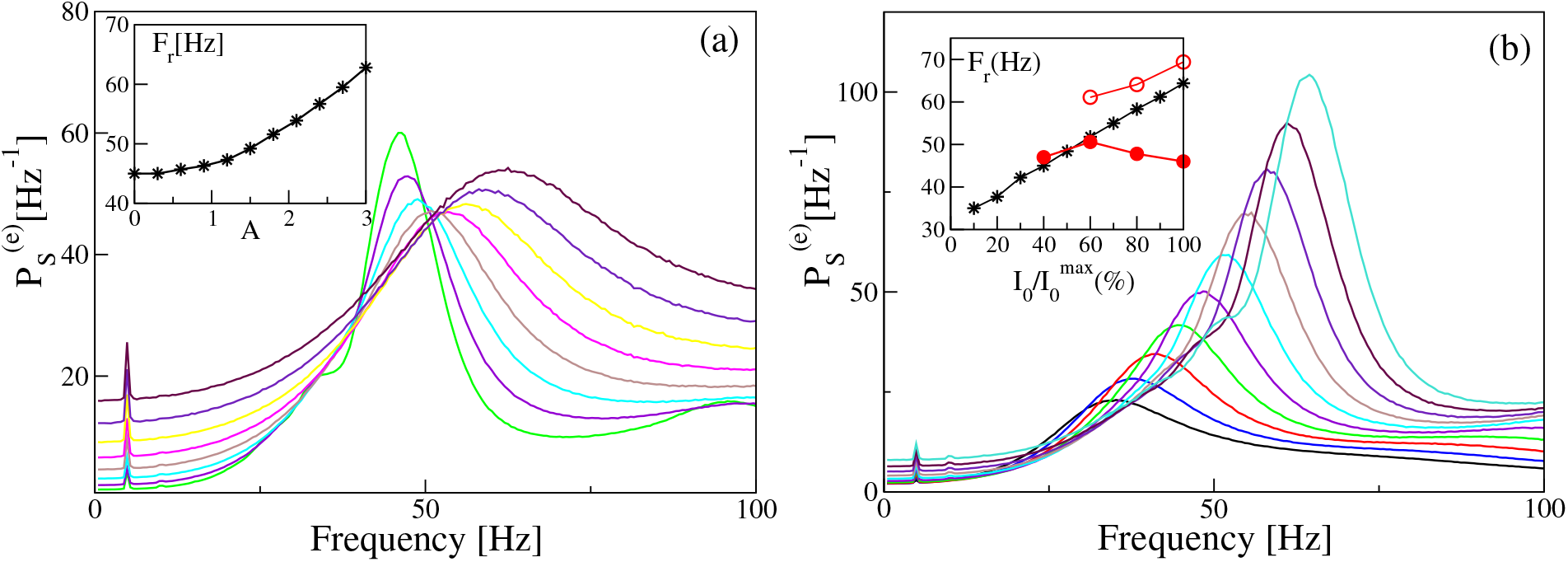
Power spectra dependency on noise and forcing amplitudes (PING set-up) Power spectra 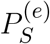 for different noise level *A* (a) and different amplitude of the external input *I*_0_ (b), for a fixed forcing frequency *ν_θ_* = 5 Hz. In the insets are reported the frequencies *F_r_* of the main peak as a function of the noise level (a) and of the amplitude of the external drive *I*_0_ (b). In the inset of panel (b) are also reported experimental data extracted from Fig. 4F of Ref (Butler et al., 2016) (filled red circles) and from Fig. 4D of Ref. (Butler et al., 2018) (open red circles). The curves in (a) are obtained by varying the noise amplitude *A* ∈ [0.9 : 3.0] with a step of 0.3, while keeping fixed *I*_0_ = 10. On the other hand the curves in (b) refer to different forcing amplitudes 2 ≤ *I*_0_ ≤ 20, varied in steps of 0.2, with fixed noise amplitude *A* = 1.4. The other parameters are as in Fig. 4.

From the previous analysis we have understood that, for constant forcing frequency, the *γ*-peak shifts towards higher frequencies by increasing the forcing amplitude or the noise level.

Therefore to obtain an increase of *F_r_* with the forcing frequency *ν_θ_*, analogously to the results reported in (Butler et al., 2016) (and displayed as filled red circles in Fig. 10(a) and (c)), we should perform numerical experiments where *ν_θ_* increases together with *A* or *I*_0_. The simplest protocol is to assume that *A* (*I*_0_) will increase linearly with *ν_θ_*. The results obtained for the PING (ING) set-up are reported in Fig. 12 (a) (Fig. 12 (b)). As evident from the figures, in both set-ups and for both protocols we obtain results in reasonable agreement with the experiment. In the present framework, we have also analyzed the dependence of the *γ*-power *P_γ_* on *ν_θ_*. In particular this quantity increases almost linearly with the forcing frequency, at variance with the experimental results in (Butler et al., 2016) which revealed a sort of resonance with an associated maximal *γ*-power around *ν_θ_* = 5 Hz (the experimental data are displayed as red circles in the insets of Fig. 10(b) and (d)).

**Figure 12.**
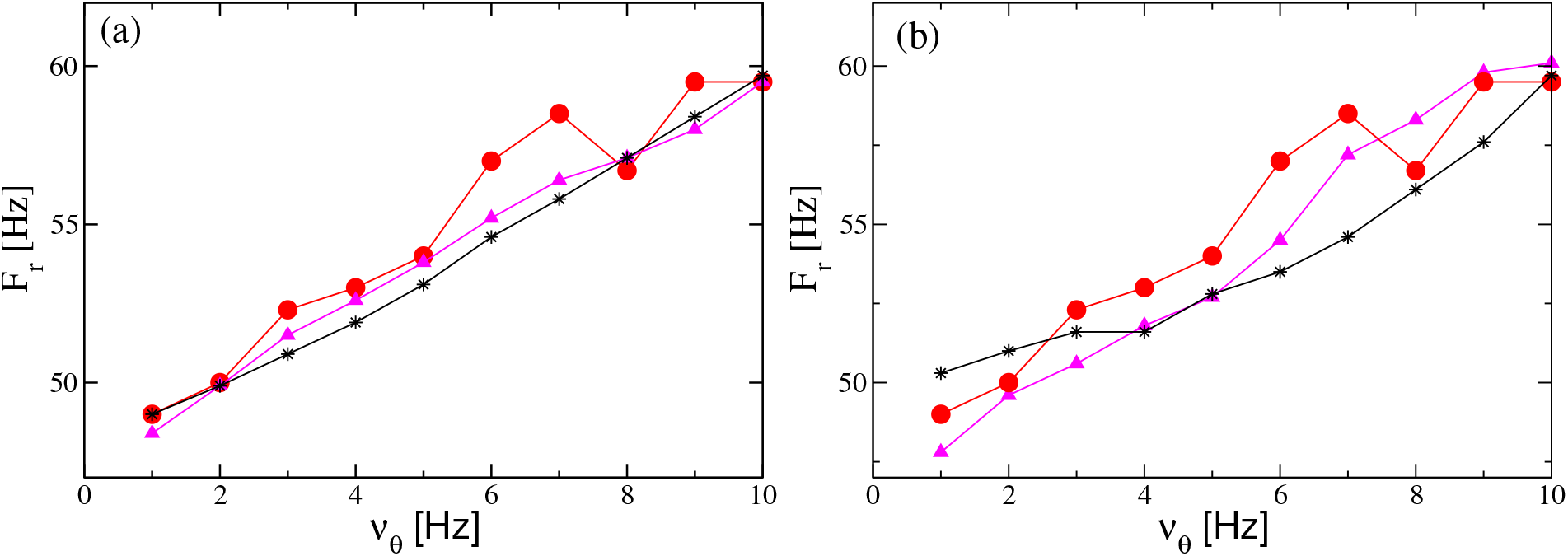
Influence of the theta frequency on the gamma oscillations. Frequency *F_r_* of the main peak of the power spectrum 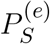 versus *ν_θ_* for the PING (a) and ING (b) set-ups. Red filled circles represent the experimental data extrapolated from Fig. 4C in (Butler et al., 2016). Black stars (magenta triangles) refer to numerical data obtained by varying linearly the noise amplitude *A* (the forcing amplitude *I*_0_) as a function of *ν_θ_* and maintaining the forcing amplitude *I*_0_ (the noise amplitude *A*) constant. The data shown as black stars for the PING (ING) set-up in panel (a) (panel (b)) are obtained by adding white noise to the evolution of the mean membrane potentials and by varying linearly its amplitude in the interval *A* ∈ [1.4 : 2.9] as a function of *ν_θ_* with *I*_0_ = 10 (*I*_0_ = 9). The magenta triangles refer to data obtained by keeping fixed the noise amplitude at the value *A* = 1.4 and by varying linearly with *ν_θ_* the forcing amplitude *I*_0_ in the range [9.5 : 18] ([8 : 14]) for the PING (ING) set-up in panel (a) (panel (b)). Other parameters for as in Fig. 4.

## 5 DISCUSSION AND CONCLUSIONS

In this paper we have analyzed the dynamics of a new class of neural mass models arranged in two different set-ups: an excitatory-inhibitory network (or PING set-up) and a purely inhibitory network (or ING set-up). The considered neural mass models are extremely relevant to mimick neural dynamics for two reasons. On one side because they are not derived heuristically, since they reproduce exactly the dynamics of excitatory and inhibitory networks of spiking neurons for any degree of synchronization (Montbrió et al., 2015; Devalle et al., 2017; Ceni et al., 2019). On another side these neural masses reproduce the macroscopic dynamics of quadratic integrate-and-fire neurons, which are normal forms of class I neurons, therefore they are expected to represent the dynamics of this large class of neurons (Ermentrout and Kopell, 1986).

In this paper we have shown that *θ*-nested *γ* oscillations can emerge both in the PING and ING set-up under an external excitatory *θ*-drive whenever the system, in absence of forcing, is in a regime of asynchronous dynamics, but in proximity of a Hopf bifurcation towards collective *γ* oscillations. The external forcing drives the system across the bifurcation inside the oscillatory regime, thus leading to the emergence of *γ* oscillations. The amplitude of these collective oscillations is related to the distance from the bifurcation point, therefore it depends on the phase of the *θ*-forcing term. These nested oscillations can arise both in proximity of super-critical and sub-critical Hopf bifurcations. However, in the latter case their amplitudes are no more symmetric with respect to the maximum value of the theta stimulation, somehow analogously to the experimental findings reported in (Butler et al., 2016).

Analogous results have been reported for an excitatory-inhibitory network with a recurrent coupling among the excitatory neurons, by considering the Wilson-Cowan rate model (Onslow et al., 2014). However, at variance with our neural mass model, the Wilson-Cowan one fails to reproduce the emergence of *γ*-oscillations, displayed by the corresponding spiking networks, in several other set-ups. In particular, the Wilson-Cowan model is unable to display COs for purely inhibitory populations (the ING set-up), without the addition of a delay in the IPSPs transmission, delay that is not required in the network model. Moreover the Wilson-Cowan model is unable to display COs even for excitatory-inhibitory coupled populations in absence of a recurrent excitation (Onslow et al., 2014; Devalle et al., 2017). As shown in Appendix B, the considered neural mass model in the PING set-up displays clear *θ*-nested *γ* oscillations in absence of any recurrent coupling or with recurrent couplings only among the inhibitory neurons.

Furthermore, we have identified two different types of phase amplitude couplings. One characterized by a perfect locking between *θ* and *γ* rhythms, corresponding to an overall periodic behaviour dictated by the slow forcing. The other one where the locking is imperfect and dynamics is quasi-periodic or even chaotic. The perfectly locked *θ*-nested *γ* oscillations display at the same time two types of cross-frequency coupling: phase-phase and phase amplitude coupling (Hyafil et al., 2015). These states arise for *ν_θ_* larger than 2-3 Hz and for sufficiently large forcing amplitudes. From the results reported in (Butler et al., 2016) for the CA1 region of the hippocampus under sinusoidal forcing *in vitro*, it is evident that perfectly phase locked PACs have been observed in each single slice. However, *in vivo* this perfect phase-phase locking cannot be expected, see the detailed discussion of phase-phase coupling reported in (Scheffer-Teixeira and Tort, 2016), where the authors clarify that phase locking is indeed observable, but only over a limited number of successive *θ*-cycles. Therefore, PAC with an underlying chaotic (or noisy) dynamics is the scenario usually expected in behaving animals.

From our analysis it emerges also that locked states are more frequent in the ING set-up. The purely inhibitory population is more easily entrained by the forcing with respect to the coupled excitatory-inhibitory population system, where the forcing is applied to the excitatory population. This result is somehow in agreement with recent findings based on the analysis of phase response curves, which suggest that stimulating the inhibitory population facilitates the entrainment of the gamma-bands with an almost resonant frequency (Akao et al., 2018; Dumont and Gutkin, 2019). However, these analyses do not consider *θ*-*γ* entrainment: this will be a subject of future studies based on exact macroscopic phase response curves (Dumont et al., 2017; Dumont and Gutkin, 2019).

Our modelization of the PAC mechanism induced by an external *θ*-forcing is able to reproduce several experimentals features reported for optogenetic experiments concerning the region CA1, CA3 of the hippocampus, as well as MEC (Akam et al., 2012; Pastoll et al., 2013; Butler et al., 2016, 2018). In agreement with the experiments, we observe nested *γ* COs for forcing frequencies in the range [1 : 10] Hz, whose amplitude grows proportionally to the forcing one. Furthermore, the *γ* power and the frequency of the *γ*-power maximum increase almost linearly with the forcing amplitude. However, the neural mass model in all the examined PING and ING set-ups is unable to reproduce the increase in frequency of the *γ*-power peak with *ν_θ_* reported in (Butler et al., 2016). Indeed such effect was expected by the observation that during movement, both the frequencies of hippocampal theta oscillations (Sławińska and Kasicki, 1998) and gamma oscillations (Ahmed and Mehta, 2012) increase with the running speed of the animal. However, the variation of the *γ* frequency reported in (Ahmed and Mehta, 2012) for behaving animals amounts almost to 100 Hz, while in the optogenetic experiment by Butler et al. (Butler et al., 2016), the increase was limited to ≃ 10 Hz. In order to get a similar increase in the neural mass model, we have been obliged to assume that the noise (or the forcing amplitude) increases proportionally to *ν_θ_*. On one side, further experiments are required to clarify if, during optogenetic experiments, the forcing (or noise amplitude) affecting the neural dynamics is indeed dependent on *ν_θ_*. This could be due to a reinforcement of the synaptic strenghts for increasing forcing frequencies, or to the fact that higher *θ* frequencies can favour neural discharges in regions different from CA1, that can be assimilated to external noise. On another side it should be analysed if other bifurcation mechanisms, beside the Hopf one, here considered, can give rise to such a dependence of *γ* power on *θ* forcing.

## FUNDING

AT received financial support by the Excellence Initiative I-Site Paris Seine (Grant No ANR-16-IDEX-008) (together with HB and MS), by the Labex MME-DII (Grant No ANR-11-LBX-0023-01) (together with SO) and by the ANR Project ERMUNDY (Grant No ANR-18-CE37-0014), all part of the French programme “Investissements d’Avenir”.

## ACKNOWLEDGMENTS

This work has been initiated during the Advanced Study Group 2016/17 “From Microscopic to Collective Dynamics in Neural Circuits” at the Max-Planck Institute for Physics of Complex System in Dresden (Germany), where SO and AT had extremely useful interactions with E. Montbrió and H.P. Robinson. We also acknowledge fruitful discussions with D. Angulo-Garcia, F. Devalle, G. Dumont, M. di Volo, A. Pikovsky.

## AUTHOR CONTRIBUTIONS

MS, HB, and SO performed the simulations and data analysis. AT was responsible for the state-of-the-art review and the paper write-up. All the authors conceived and planned the research.

## CONFLICT OF INTEREST STATEMENT

The authors declare that the research was conducted in the absence of any commercial or financial relationships that could be construed as a potential conflict of interest.

## APPENDIX A: PING SET-UP: SUB-CRITICAL HOPF

In the PING set-up with only recurrent excitation (i.e. with *J*^(*ee*)^ ≠ 0, *J*^(*ii*)^ = 0 and *J*^(*ie*)^ = *J*^(*ei*)^ ≠ 0), it is possible to observe the emergence of COs also via a sub-critical Hopf bifurcation, by using *H*^(*e*)^ as control parameter, as shown in Fig. 13 (a). This is due to the nature of the Hopf bifurcation that can be modified by simply varying the value of *H*^(*i*)^. In this case we observe three regimes: an asynchronous one for 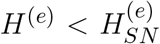; an oscillatory one for 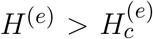 and a bistable one in the range 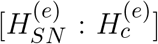. The frequency of the COs *ν*^(*e*)^ is always in the *γ* range with a minimal value ≃ 36 Hz achievable at the Hopf bifurcation, see the inset of Fig. 13 (a).

**Figure 13.**
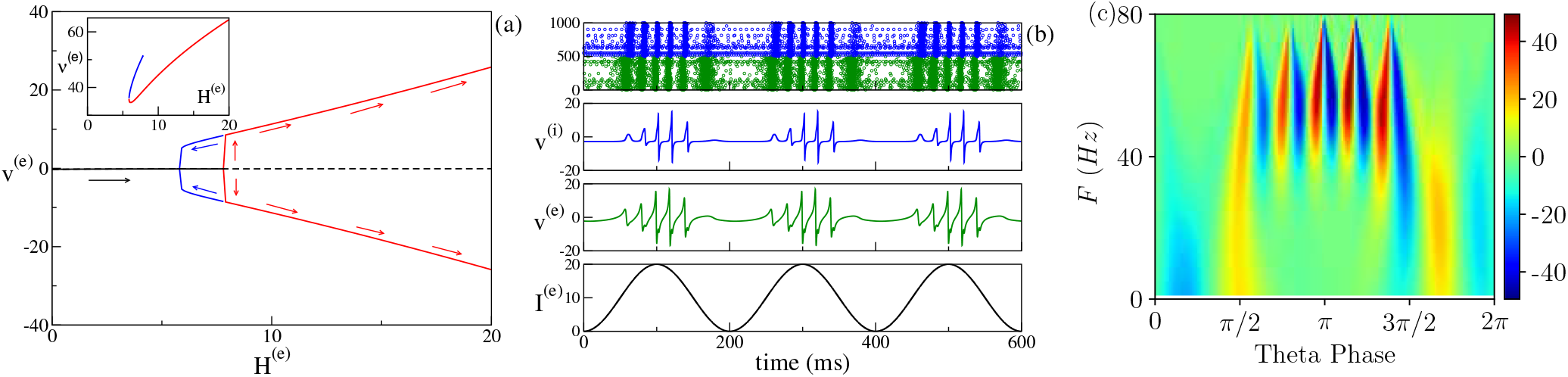
(PING set-up: subcritical Hopf) (a) Bifurcation diagram of the average membrane potential *υ*^(*e*)^ as a function of *H*^(*e*)^. The black continuous (dashed) line identifies the stable (unstable) fixed point. The red lines denote the extrema of the limit cycles. The subcritical Hopf bifurcation occurs at 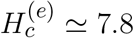 while the saddle-node of limit cycles at 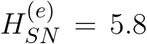. In the inset the COs’ frequency *ν*^(*e*)^ is displayed as a function of *H*^(*e*)^. (b) From top to bottom: raster plot where green (blue) dots refer to excitatory (inhibitory) neurons; average membrane potentials *υ*^(*i*)^ and *υ*^(*e*)^ as obtained by the evolution of the neural mass models and forcing current *I*^(*e*)^ for 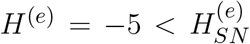 and *ν_θ_* = 5 Hz. (c) Continuous wavelet transform over a single *θ* cycles for *υ*^(*e*)^ with system setting as in (b). The remaining system parameters are *J*^(*ee*)^ = 8, *J*^(*ii*)^ = 0, *J*^(*ie*)^ = *J*^(*ei*)^ = 10, *H*^(*i*)^ = −8.0

If we consider the unforced system with 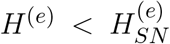 and we apply a *θ*-forcing, we observe PAC oscillations. However when considering *υ*^(*e*)^, the COs are now asymmetric with respect to the maximum of the stimulation current *I*^(*e*)^ = *I_θ_*(*t*) (see Fig. 13 (b)). This effect is even more pronounced by observing the wavelet spectrogram reported in Fig. 13 (c), where a clear PFC is also observable. The asymmetry in the onset of the gamma oscillations is clearly visible in the continuous wavelet transform obtained from the experimental data and reported in Fig. 4G in (Butler et al., 2016). This asymmetry can be explained in an adiabatic framework by considering the corresponding bifurcation diagram shown in Fig. 13 (a). Indeed for the sub-critical Hopf, the COs will emerge for 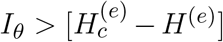, but they will disappear for a different value of the forcing, namely 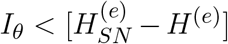. Instead for a super-critical Hopf the emergence and disappearence of the oscillations will occur at the same forcing amplitude, namely 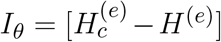.

## APPENDIX B: DIFFERENT PING SET-UPS

In the article we have considered an unique configuration giving rise to COs via the PING mechanism: namely, two cross coupled inhibitory and excitatory populations with recurrent excitation and no recurrent inhibition (i.e. *J*^(*ee*)^ ≠ 0 and *J*^(*ii*)^ = 0). However, other network configurations can give rise to PING induced oscillatory regimes. In particular, we have observed such oscillations with only cross-couplings in absence of recurrent excitation and inhibition (i.e. *J*^(*ee*)^ = *J*^(*ii*)^ = 0), as well as in presence of recurrent inhibition only (i.e. *J*^(*ee*)^ = 0 and *J*^(*ii*)^ ≠ 0). In the following we refer to the former configuration as PING_0_ set-up, while the latter configuration with recurrent inhibition is identified as PING_*I*_ set-up. In both configurations the neural mass reproduces the emergence of *γ* oscillations via a super-critical Hopf bifurcation for increasing values of *H*^(*e*)^, as shown in Figs. 14 (a) and (b). Indeed the frequencies of the COs are in the range [26 : 63.5] Hz ([29.1 : 53.9] Hz) for PING_0_ (PING_*I*_) set-up. In both configurations the corresponding bifurcation, as a function of the parameter *H*^(*i*)^, is sub-critical and COs disappear for sufficiently positive values of *H*^(*i*)^, analogously to what reported in the main text for the PING set-up with only recurrent excitation. It should be stressed that the standard Wilson-Cowan neural mass model gives rise to COs only in presence of a recurrent excitation (Wilson and Cowan, 1972), thus being unable to reproduce the spiking network dynamics (Dumont and Gutkin, 2019).

**Figure 14.**
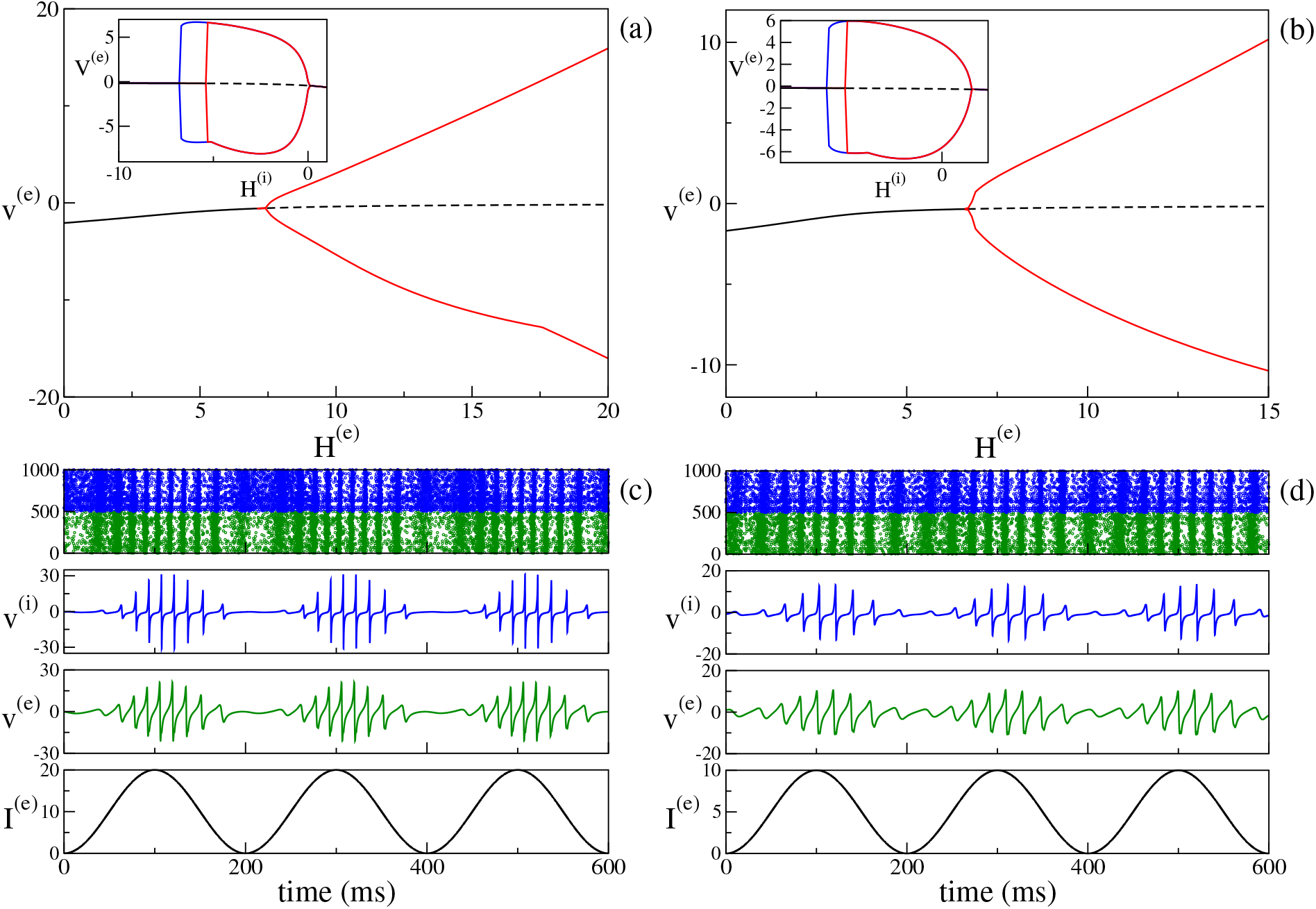
(Different PING set-ups) Bifurcation diagram versus *H*^(*e*)^ for the PING_0_ (a) and PING_*I*_ (b) set-ups for *H*^(*i*)^ = −0.5. In the corresponding insets, are reported the bifurcation diagrams as a function of *H*^(*i*)^, for *H*^(*e*)^ = 10. *θ*-nested *γ* oscillations emerging in the PING_0_ (c) and PING_*I*_ (d) configurations for *I*_0_ = 20 and *ν_θ_* = 5 Hz. From top to bottom the raster plot where green (blue) dots refer to excitatory (inhibitory) neurons; the average membrane potentials *υ*^(*i*)^ and *υ*^(*e*)^ as obtained by the evolution of the neural mass models and the forcing currents *I*^(*e*)^. Parameters for the PING_0_ set-up are *J_ee_* = *J_ii_* = 0, while for PING_*I*_ are *J_ee_* = 0 and *J_ii_* = 8. In both cases *J_ie_* = *J_ei_* = 10 and *H*^(*i*)^ = −0.5. In the corresponding insets we set *H*^(*e*)^ = 10.

In presence of an external *θ*-forcing with *ν_θ_* = 5 Hz, we clearly observe *θ*-nested *γ* oscillations, as shown in the raster plots reported in Figs. 14 (top rows of panels (b) and (d)). These oscillations are phase amplitude modulated from the forcing, as it results to be evident from the shape of the mean membrane potentials *V*^(*e*)^ and *V*^(*i*)^ reported in the middle rows of Figs. 14 (c) and (d).

